# Linking microscopic and mean-field descriptions of dopaminergic modulation in neural circuits

**DOI:** 10.1101/2025.09.01.673498

**Authors:** Gabriele Casagrande, Augustinas Fedaravičius, Chloe Duprat, Anthony Randal McIntosh, Pierpaolo Sorrentino, Spase Petkoski, Ausra Saudargiene, Viktor Jirsa, Damien Depannemaecker

**Affiliations:** Aix Marseille Universitè, INSERM, Institut de Neurosciences des Systèmes, Marseille, France; Neuroscience Institute, Lithuanian University of Health Sciences, Kaunas, Lithuania; Department of Informatics, Vytautas Magnus University, Kaunas, Lithuania; Simon Fraser University, Vancouver, British Columbia, Canada

## Abstract

Next-generation neural mass models capture the collective behavior of brain regions through an exact mean-field reduction of the spiking neural networks, retaining key microscopic properties while remaining computationally efficient. This explicit link to single-neuron dynamics makes them a natural framework for studying how neuromodulators, such as dopamine (DA), shape macroscopic network activity. Integrating its effect is a key step toward biologically grounded brain models. Because the spiking network underlying such a reduction is fully known, the reduced model can be directly validated against it. Here, we perform a systematic comparison between a recently proposed DA-modulated mean-field model and its corresponding spiking network across multiple dynamical regimes, evaluating its ability to capture the qualitative organization of the network activity, and transitions between dynamical states. This comparison identifies the conditions under which the reduction accurately reflects the behavior of the full network, and highlights its theoretical limitations.

## 1 Introduction

The brain, like many complex biological systems, consists of a vast number of microscopic units. Their interactions shape the emergent collective behaviors that characterize brain function, making the microscopic dynamics of individual neurons and their connections a natural starting point for understanding brain-wide phenomena. However, the majority of human brain recordings used to analyze these dynamics (e.g., fMRI, PET, EEG) are captured at the mesoscopic scale, reflecting the aggregated activity of thousands of neurons. To model this specific scale, Neural Mass Models (NMMs) have been extensively developed, offering varying degrees of biological accuracy [1] ranging from phenomenological to biophysically grounded formulations [2, 3, 4, 5, 6]. To bridge between the micro and mesoscopic scales, mean-field techniques have been employed to reduce the full spiking network description into collective macroscopic observables, such as the population firing rate and mean membrane potential. Among these, next-generation NMMs have played a particularly prominent role. Initially developed by Montbrió et al. [7] for globally coupled networks of quadratic integrate-and-fire (QIF) neurons, this framework combines the Ott-

Antonsen ansatz with a Lorentzian assumption on the distribution of neuronal parameters to obtain a closed low-dimensional set of ordinary differential equations for the population firing rate and mean membrane potential in the thermodynamic limit. This formulation is particularly powerful because it yields a computationally and analytically efficient reduction while maintaining an exact mathematical link to the underlying microscopic system. Subsequently, it has been expanded to accommodate more complex single-neuron mechanisms, such as adaptation [8, 9], electrical synapses [10], and delays [11], and to reproduce higher-level cognitive phenomena such as working memory [12, 13]. Among these extensions, particular attention has been devoted to capturing the effect of neuromodulation [6, 14], the mechanisms by which neurotransmitters shape neuronal excitability and network-level activity [15]. Together with anatomical connectivity, these substances enable the brain’s rich repertoire of activities [16, 17, 18]. Among them, dopamine (DA) is particularly important. DA is heavily involved in fundamental processes ranging from motor control [19, 20, 21] to learning and the processing of rewarding stimuli [22, 23, 24]. Consequently, dysfunctions in dopaminergic pathways are a hallmark of pathological conditions such as schizophrenia and Parkinson’s disease [25]. Despite the broad adoption of these mean-field formulations, most studies focus primarily on the mathematical derivation of the mean-field equations or on their subsequent analysis, treating the microscopic network mainly as a starting point rather than as a benchmark against which the reduction is validated. Hence, systematic comparisons between the reduced mesoscopic description and the underlying microscopic network dynamics remain rarely explored [9]. As a consequence, it remains unclear exactly which aspects of network dynamics are preserved by the mean-field reduction, where the approximation breaks down, and how these discrepancies vary with parameters such as neuromodulator concentration. Addressing this gap, the present work builds on a previously validated formulation of a specific population of interconnected neurons under DA modulation [14], further incorporating the activity of DA receptors that alter neuronal behavior in a concentration-dependent manner. Because the exact composition of the underlying population is known, the mean-field approximation can be rigorously benchmarked against the full network dynamics. We first assess the ability of the mean-field formulation to capture the collective behavior of the network and tracks the transitions between different dynamical regimes and we provide an extensive qualitative analysis of the mismatch between the two formulations.

Finally, having established the mean-field model capacity to capture the network underlying dynamics, we explore whether the reduced model preserves more complex features suitable for direct comparison with experimental data. Ultimately, this work proposes a comprehensive mathematical framework to investigate the modulatory role of DA in shaping neural population dynamics. By formally clarifying the connection between single-neuron electrophysiology and the network-level mesoscopic effects, this framework aims to provide a robust theoretical basis for interpreting empirical data. Similar integrative approaches have already proven valuable for exploring personalized treatment strategies in other neurological disorders, such as epilepsy [26], and have begun to be applied to DA-dependent pathologies such as Parkinson’s disease and schizophrenia within whole-brain modeling frameworks [27, 28]. In this context, by clarifying where and how the mean-field reduction deviates from the underlying microscopic dynamics, the present work may help guide the identification of suitable dynamical regimes and the exploration of parameter space in these whole-brain models, ultimately supporting their clinical applicability.

## 2 Results

### 2.1 Dynamical regimes of the mean-field reduction

We find that the reduced model effectively captures the behavior of the spiking neural network across several dynamical regimes.

By varying the parameter *u*_*jump*_, i.e. the after-spike jump in the adaptation variable, we observe the emergence of three distinct dynamical regimes, represented in Fig. 1, namely asynchronous spiking, sustained oscillations, and population bursting. We point out that the asynchronous state can occur in two mutually exclusive forms, in separate region of the parameter space explored: one characterized by a high level of activity, in which most neurons are firing (shown in Fig. 1a), and the other characterized by a low level of activity, in which only a reduced subset of neurons (those associated to higher values of the heterogeneity current) are active.

**Figure 1:**
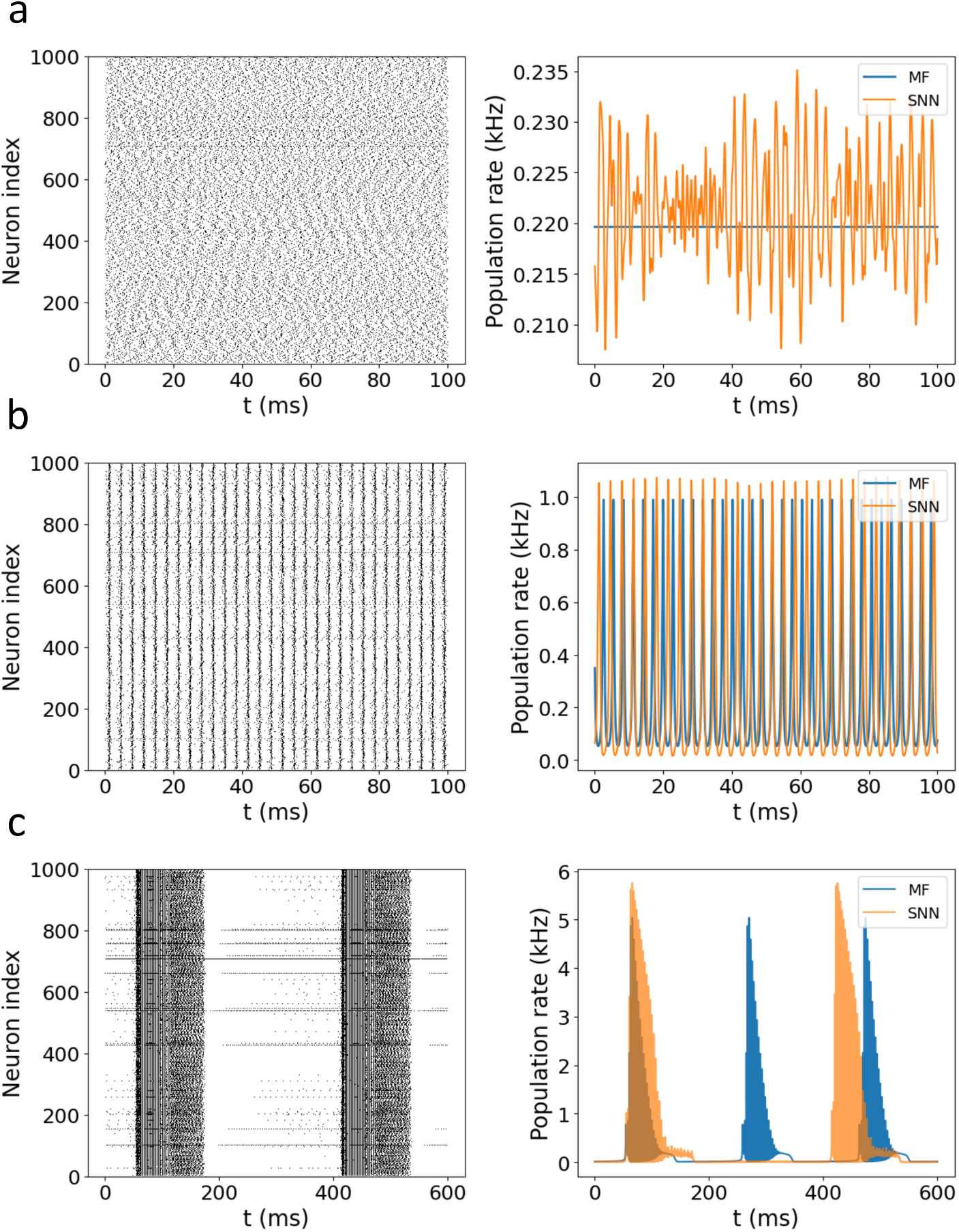
Dynamical regimes of mean-field reduction compared to spiking neural network. Panels on the left for each row represent the raster plot of 1000 randomly selected neurons inside the population of size *N* = 5000. Each black dot represents the time in which the corresponding neuron fires a spike. On the right, time series of the population firing rate, both for the mean-field model (blue) and the spiking neural network (orange) are represented. By adapting the parameter *u*_*jump*_ different dynamical regimes are achieved. In particular, we can distinguish: (a) Asynchronous spiking (*u*_*jump*_ = 9.25), where individual neurons fire regularly, but without any synchronization with the others. In the mean-field model, the constant population rate is consistent with convergence to a stable equilibrium. (b) Sustained oscillations (*u*_*jump*_ = 5.0), characterized by fast periodic activity of the population firing rate and synchronous spiking behavior at network level. (c) Population bursting (*u*_*jump*_ = 10.25), characterized by intermittent intervals of fast synchronous activity within the network (active phase) and near quiescent periods (silent phase).

We note that, near the boundary between asynchronous and population bursting, the two dynamical regimes can coexist under the same parameter values, depending on the initial condition.

In the asynchronous regime (Fig. 1a), the mean-field prediction for the population rate is constant, while the spiking network exhibits visible fluctuations around this mean value. This reflects the essence of the mean-field formalism, naming the ability to describe the behavior of a large neuronal ensemble, through a reduced set of collective variables which describe the overall averaged behavior of the neural population.

Nevertheless, we also observe a quantitative discrepancy between the two descriptions. This discrepancy is most evident in the oscillatory regimes, namely sustained oscillations (Fig. 1b) and population bursting (Fig. 1c). In both cases, the characteristic frequencies of the spiking network and of the mean-field model do not match. Oscillations and bursts have comparable waveforms and amplitudes in the two descriptions, but their timing differs.

Quantitative discrepancies between the mean-field and spiking-network descriptions have also been reported in several previous studies [9]. These discrepancies have been associated with violations of the assumptions used in the reduction, including strong spike-frequency adaptation, finite spike-reset conditions, and strong neuronal heterogeneity. Importantly, although these effects can shift the bifurcation parameters, the qualitative bifurcation structure may remain preserved.

### 2.2 Transition between dynamical regimes

Beyond verifying the correspondence between the reduced model and the underlying network, the mean-field formalism also makes it possible to understand how the emergence of different macroscopic behaviors is linked to microscopic mechanisms. This allows us to assess which aspects of the microscopic dynamics are captured by the mean-field description, and which are most relevant for achieving good correspondence between the two formalisms.

We first focus on the transition between sustained oscillations and the asynchronous state, depicted in Fig. 2. From the network perspective, this transition is characterized by an increasing level of asynchrony among neurons. This is illustrated in Fig. 2a-b, where histograms of the coefficient of variation CV [29, 30] and of CV2 [31, 32], are shown together with the Kuramoto order parameter *R* [33]. These three measures provide insight into the underlying microscopic mechanisms of the transition.

**Figure 2:**
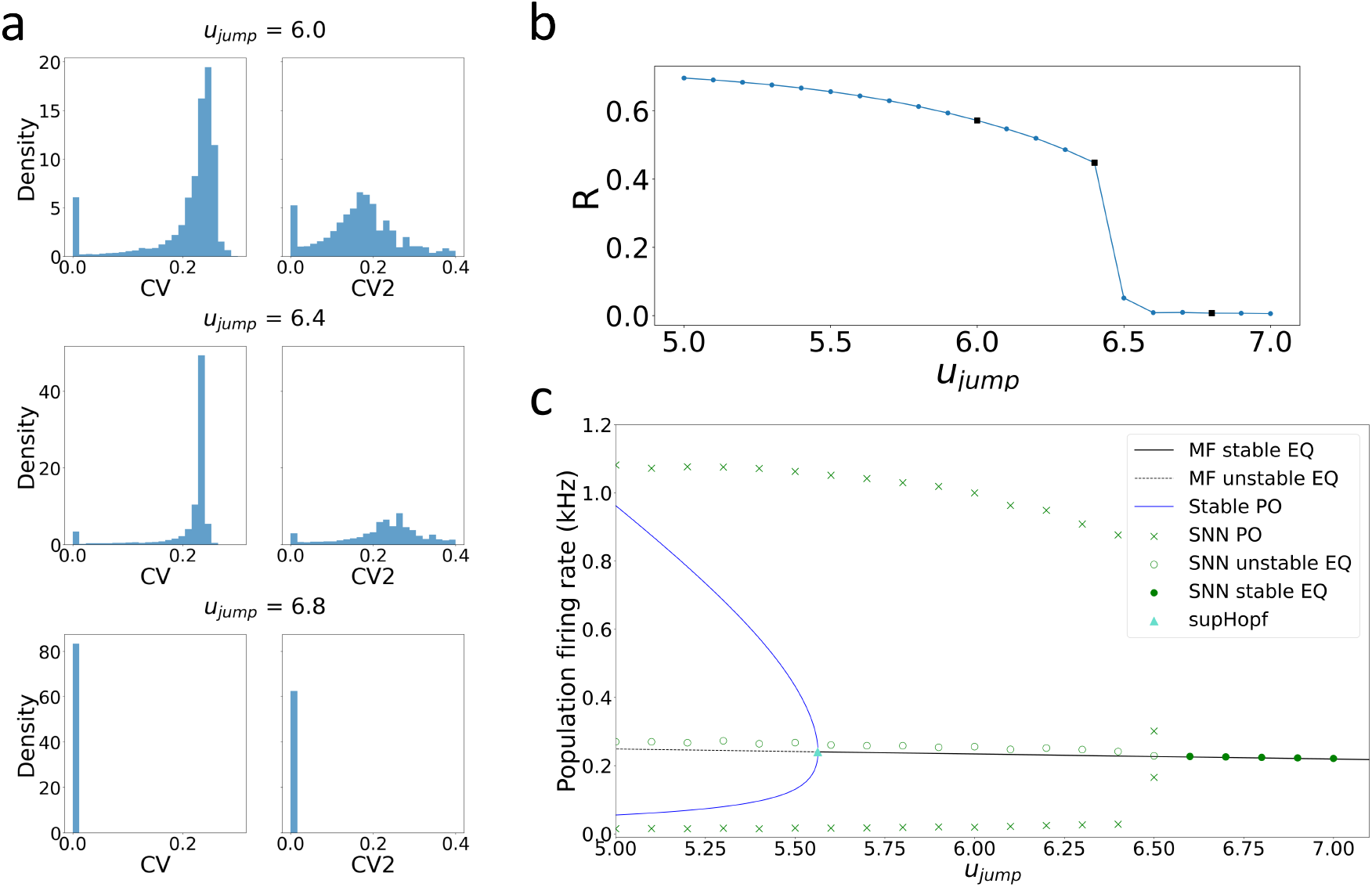
Detailed view of the transition between sustained oscillations and the asynchronous regular regime. (a) Distributions of CV and CV2 across the population of *N* = 5000 neurons, shown at three representative values of *u*_*jump*_ spanning the transition. In the sustained oscillation regime, both measures are distributed around values below 1 (*CV* ≈ 0.22, *CV* 2 ≈ 0.18), reflecting globally and locally regular firing. After the transition to the asynchronous regular state (*u*_*jump*_ = 6.8), both distributions collapse toward 0, indicating a further increase in spiking regularity. (b) Kuramoto order parameter *R* as a function of *u*_*jump*_. *R* decreases toward the critical value of the bifurcation parameter and is close to 0 past the transition, confirming the loss of global synchrony between neurons within the network at the transition. Marked black squares represent the values of *u*_*jump*_ for which the distribution in (a) were presented. (c) Comparison between the mean-field and spiking-network descriptions in the region of interest. Black lines represent the model stable (solid) and unstable (dashed) equilibrium, while the blue curves show the extremes of the periodic orbits. Green circles represent instead the equilibrium of the spiking neural network (filled for the stable equilibrium and empty for the unstable one). Finally, green crosses indicate the maximum and minimum population firing rates measured from the spiking-network oscillations. The transition occurs at different values of *u*_*jump*_ in the two formulations: in the spiking network it is located at 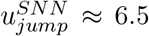, while in the mean field it corresponds to a supercritical Andronov-Hopf bifurcation at 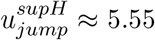. Despite this quantitative offset, the underlying mechanism is qualitatively similar in both descriptions, with oscillations of decreasing amplitude collapsing onto a fixed-point equilibrium. However, while in the mean-field reduction amplitude of oscillations gradually approach the fixed point, in the network the transition is comparatively sharper.

At *u*_*jump*_ = 6.0, both CV and CV2 are concentrated at low values, reflecting the global and local regularity typical of synchronous oscillations. This is further supported by the correspondingly high value of *R*. As *u*_*jump*_ increases toward the transition, neurons progressively desynchronize, as it is shown by the decreasing value of the Kuramoto order parameter. Beyond the transition (*u*_*jump*_ = 6.8), synchronization is essentially absent. In contrast, the CV and CV2 distributions remain concentrated at low values, showing that individual neurons continue to fire regularly even after the population has become asynchronous. Low values of both CV and CV2 indicate that, despite the loss of synchrony, individual neurons continue to fire with high regularity, the signature of an asynchronous regular state. For clarity, raster plots illustrating this transition are reported in the Supplementary Material–Section A.

Comparing the network and mean-field descriptions (Fig. 2c), we find that the model partially captures the qualitative nature of the transition. In both descriptions, the oscillation amplitude progressively decreases as the transition is approached. In the mean-field model, this process culminates in a supercritical Andronov–Hopf bifurcation (supH) at 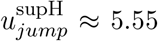, where the stable limit cycle decreases continuously in amplitude and collapses onto the stable equilibrium. The network dynamics, however, exhibit a qualitatively different loss of synchrony. Although the oscillation amplitude also decreases as the bifurcation is approached, the transition is ultimately marked by a sharp loss of synchronization rather than by the gradual collapse of the collective oscillation predicted by the mean-field model.

A residual mismatch is also observed in the precise value of the critical parameter at which the transition between the two dynamical regimes occurs. Such a discrepancy has been extensively reported in previous studies [34, 9].

A second, qualitatively distinct transition separates the asynchronous regular state from the population bursting regime (Fig. 3).

**Figure 3:**
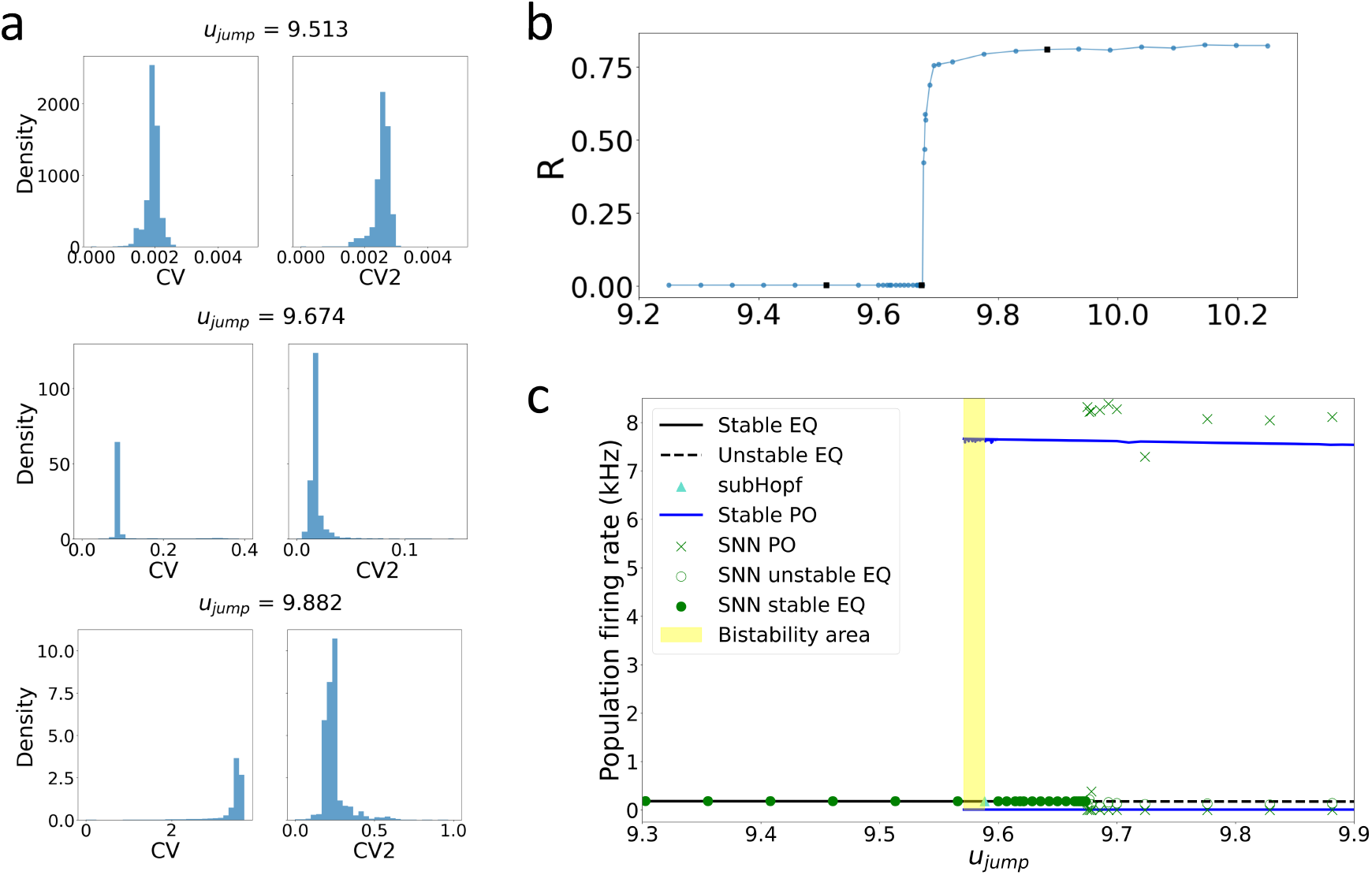
Detailed view of the transition between asynchronous regular regime and burst. (a) Distributions of the CV and of CV2, across neurons, for three values of *u*_*jump*_. Before the transition (top), at the transition (middle) and after it (bottom). Before the transition, both distributions are sharply peaked at very low values (∼ 10^−3^), indicating highly regular firing. At the transition, both shift to larger values while remaining narrow. After the transition, the CV distribution is concentrated at values above 1, whereas CV2 stays at comparatively low values (mostly *<* 0.5), consistent with regular intra-burst firing interleaved with long silent periods between population bursts. Note the different horizontal and vertical scales across panels to better represent the larga variation in the distributions observed. (b) Kuramoto order parameter *R* across the transition, showing a sharp increase in synchronization between the two regimes. Marked by black squares represent the values of *u*_*jump*_ represented in panel (a). (c) Comparison between the mean-field and spiking-network descriptions. In the model, the onset of population bursting is marked by a subcritical Andronov-Hopf bifurcation, from which an unstable limit cycle emerges. At the boundary between the two dynamical regimes there exists a region of bistability, where the system can evolve in time toward population bursting or asynchronous regime depending on its initial conditions. Within the population bursting region, the network state resides on the outer stable limit cycle, which encircles the un-stable equilibrium and the inner unstable limit cycle. Within the network the transition is similarly abrupt and, in this case, occurs in close agreement with the value predicted by the mean field. Colors of the distinct elements are the same as in the previous figure.

At the network level, the order parameter *R* jumps abruptly from values close to zero to *R* ≈ 0.8 at *u*_*jump*_ ≈ 9.67 (Fig. 3b). This is accompanied by a marked change in the CV and CV2 distributions (Fig. 3a). Below the transition (*u*_*jump*_ = 9.513), both distributions are sharply peaked at very low values (∼ 10^−3^). Close to the transition, both shift toward larger values (CV ∼ 0.1, CV2 ∼ 0.02) while remaining narrow. Beyond the transition (*u*_*jump*_ = 9.882), the CV distribution is concentrated at values higher than one, reflecting the strong global irregularity of inter-spike intervals that is characteristic of population bursting. In contrast, the CV2 distribution remains concentrated at comparatively low values (mostly *<* 0.5, with a tail approaching 1). This indicates that individual neurons fire regularly within each burst, alternating rapid spike sequences (short intra-burst intervals) with longer silent periods between population bursts. As for the previous transition, raster plots are reported in Supplementary Material, Section A.

The mean-field model reproduces this transition, which is similarly sharp and marked by a large increase in oscillation amplitude (Fig. 3c). We identify a subcritical Andronov–Hopf bifurcation at 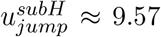, from which a branch of unstable limit cycles of increasing amplitude emerges. Because the amplitude grows rapidly over a very narrow parameter range, the periodic-orbit branch has a very steep slope with respect to *u*_*jump*_, which makes the collocation and Floquet computations poorly conditioned and prevented us from reliably following this branch by numerical continuation. Nevertheless, direct simulations of the mean-field model reveal coexistence of a stable fixed point (asynchronous state) and a stable population bursting limit cycle in a narrow bistable region starting at *u*_*jump*_ ≈ 9.57. This is consistent with the unstable branch approaching the stable upper cycle nd annihilating with it in a fold of limit cycles (FLC) [34, 35]. Note that the network transition occurs at a slightly larger value (*u*_*jump*_ ≈ 9.67) than the mean-field bistability onset.

Finally, we examine the transition between the population bursting regime and the low-activity asynchronous regime. In this specific case, the decrease in network activation makes the analysis based on the coefficient of variation and Kuramoto order parameter inadequate. Hence, to characterize this transition, we tracked the evolution of two bursting features as *u*_*jump*_ increases: the mean burst duration and the mean inter-burst interval (Fig. 4a–b). To assess the role of finite-size effects, we performed multiple simulations of the network at three different sizes spanning multiple order of magnitude (*N* = 5000, 15000, 100000), each for a duration of 30 s.

**Figure 4:**
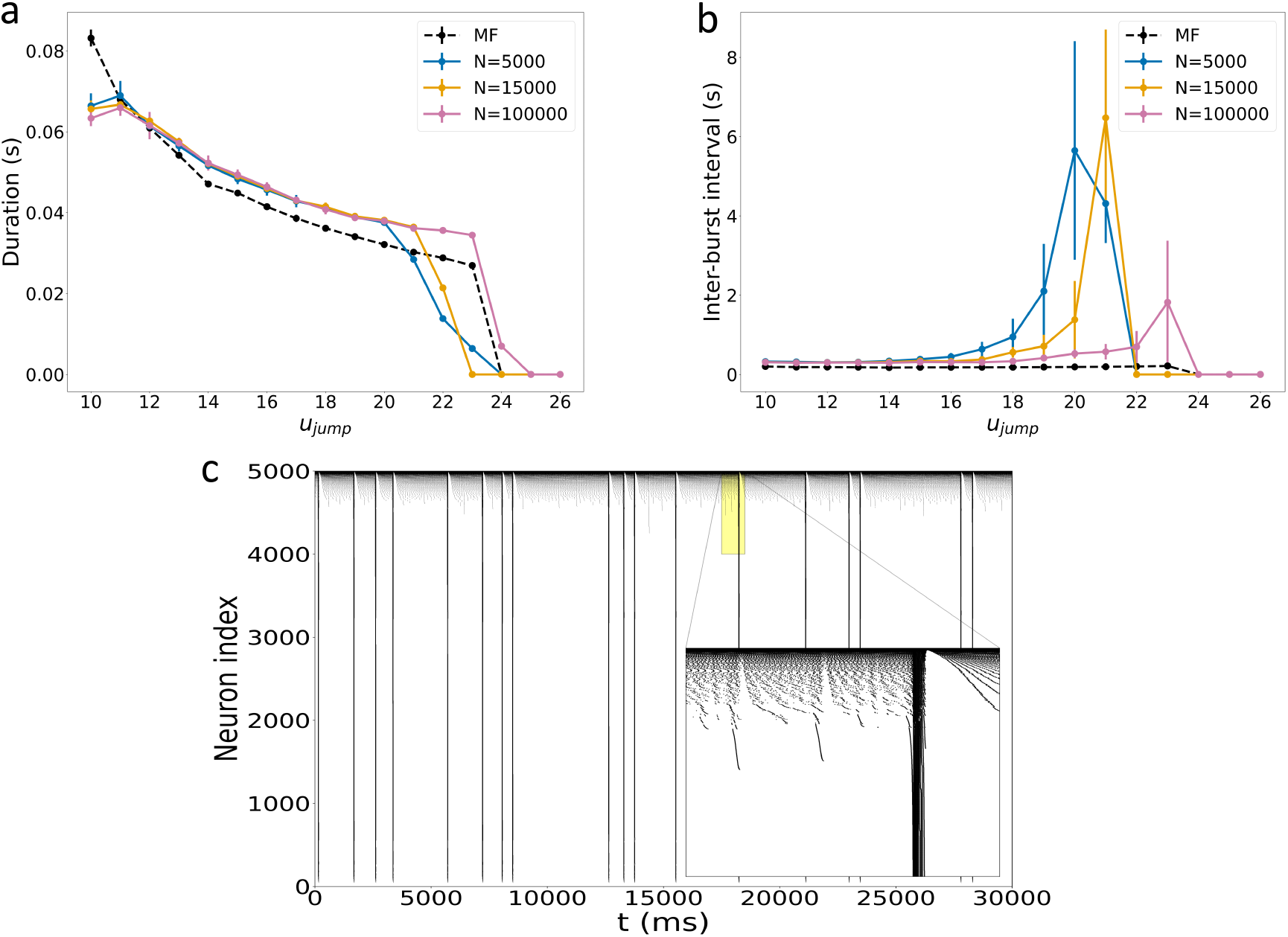
Finite-size effects on population bursting near the offset transition. (a) Mean burst duration and (b) mean inter-burst interval as a function of *u*_*jump*_, comparing the mean-field (MF, dashed black) prediction with finite networks of size *N* = 5000 (blue), 15000 (orange), and 100000 (pink). Each network simulation was performed for 30 s. For each value of the parameter and size of the network 10 simulations were realized. Error bars indicate the standard deviation of the quantity in analysis over this simulations. The mean burst duration shows a similar dependence on *u*_*jump*_ across network sizes and the MF reduction, decreasing progressively toward the offset of the population-bursting regime. In contrast, the inter-burst interval exhibits a systematic mismatch between the finite networks and the MF prediction throughout the bursting regime. While the MF value remains approximately constant, finite networks display progressively longer intervals and increasing variability as *u*_*jump*_ approaches the offset. This discrepancy reflects the increasing disrupting effect of finite-size fluctuations on the population bursting dynamics of the network, which are more evident as the firing activity of the network become sparse. The finite-size fluctuations scale approximately as 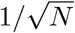, explaining the stronger deviation observed for smaller networks and the reduced discrepancy for larger N. (c) Raster plot illustrating these effects for *N* = 5000. As the offset is approached, population bursts become increasingly separated and the activity between bursts becomes more irregular, providing a direct visualization of the fluctuations emerging within the network. The inset magnifies the microscopic dynamic of the sparse fluctuations.

Both representations show a similar trend in burst duration, which decreases as the system approaches the offset of the population bursting regime, with comparable values between the two. This is not the case for the inter-burst interval. The mean-field value remains approximately constant throughout the bursting regime, whereas the network value progressively increases as the transition is approached, together with its standard deviation. This broad discrepancy reflects the lost of regularity in the inter-burst timing due to the increasing influence of finite-size effects (Fig. 4c). Near the offset, the discrepancy, which is expected to a certain extent at the transition between dynamical regimes [34, 9], is further amplified because bursting events become sparse and the low-activity state becomes increasingly sensitive to finite-size perturbations (which scale as 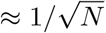). The increasing standard deviation of the network inter-burst interval therefore reflects both a reduced number of observed burst events and a genuine increase in burst-timing variability.

As for the previous transition, the offset transition is consistent with the reverse orientation of the same subcritical Andronov-Hopf/LPC structure.

### 2.3 Comparison between mean-field and spiking network dynamics

To further assess the mean-field model ability to capture the dynamical behavior of the underlying spiking network we compared the two representations over a wide range of parameters (Fig. 5). We selected two parameters of particular relevance for this purpose: the dopaminergic projection strength *c*_*dopa*_, to probe the effect of DA on network dynamics, and the after-spike increment of the adaptation variable, *u*_*jump*_.

**Figure 5:**
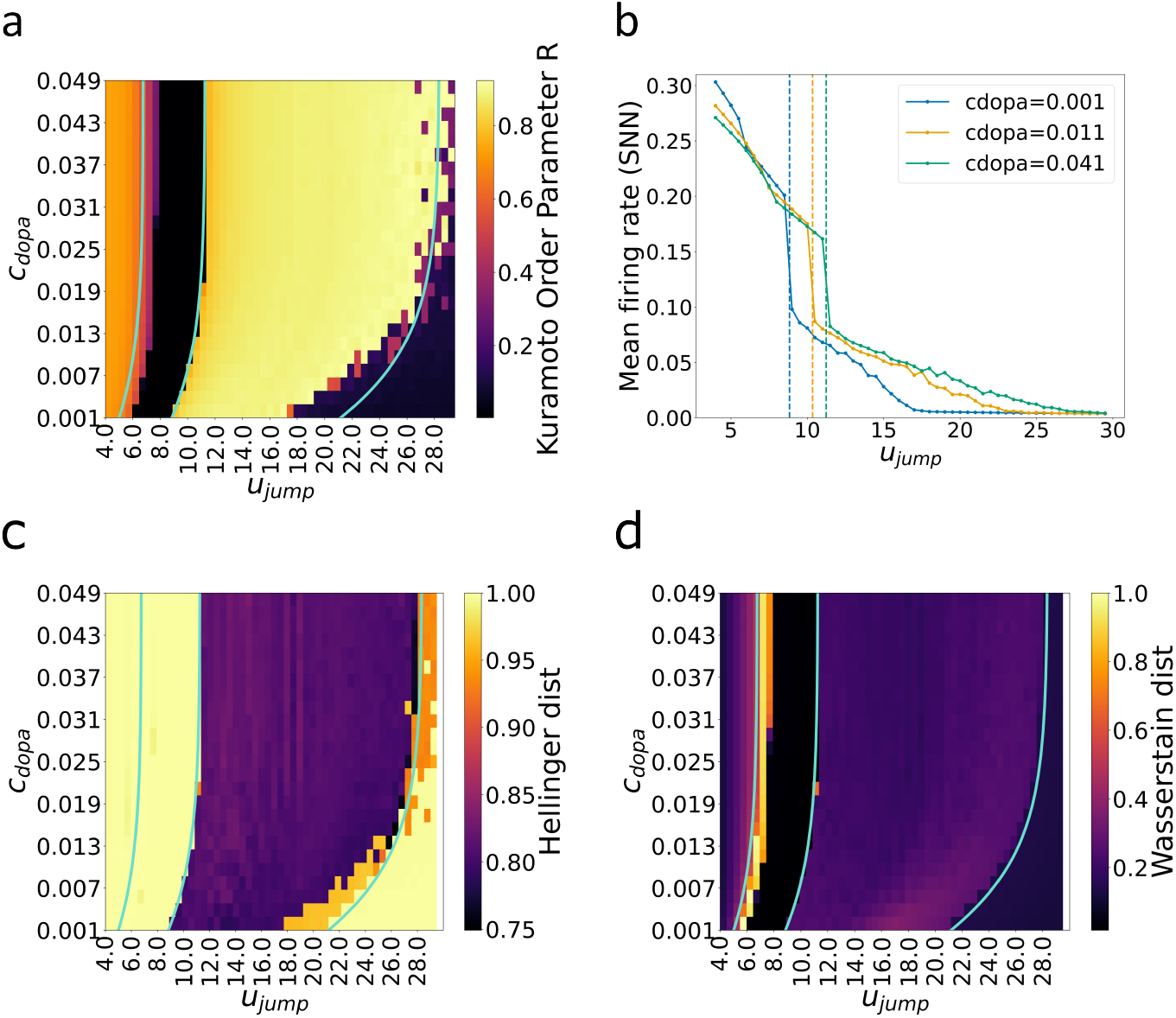
Comparison between mean-field and spiking-network dynamics across the (*u*_*jump*_, *c*_*dopa*_) parameter space. (a) Heatmap of the Kuramoto order parameter *R* for the spiking network. High values are associated with the sustained oscillation and population bursting regimes, whereas low values correspond to asynchronous regimes. Cyan curves indicate the supercritical Andronov-Hopf curve, separating the fixed-point and sustained oscillation regimes, and the subcritical Andronov-Hopf curves, delimiting the population bursting regime. The nearby FLC curves are omitted for clarity. The two representations present the same dynamical regime over a wide range of parameters, with discrepancy arising mostly at the border between distinct dynamical regimes. (b) Mean firing rate of the spiking network as a function of *u*_*jump*_ for several values of *c*_*dopa*_. Dashed vertical lines indicate the critical bifurcation values 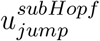 marking the onset of the population bursting regime in the mean-field. Across this transition, the spiking network exhibits a pronounced discontinuity in its mean firing rate. This facilitates the detection of the transition by the mean-field model, which describes the evolution of the mean activity but does not explicitly account for higher-order moments. In contrast, the remaining transitions are associated with much smaller changes in mean activity and are therefore less readily captured. (c) Hellinger distance across the parameter space. The largest discrepancies occur in the sustained oscillation and asynchronous regimes. In the former, the distance primarily reflects differences in the dominant oscillation frequency and spectral profile, whereas in the latter, it reflects the finite-size fluctuations associated with discrete spiking events, which are absent from the deterministic mean-field description. (d) Normalized Wasserstein distance between the mean-field and spiking network representations. The largest mismatch occurs in the transition region between the sustained oscillation and asynchronous regular regimes. Elsewhere, the distance remains comparatively low, indicating overall agreement in long-term statistical properties such as the mean activity and the distribution of oscillation amplitudes.

For the qualitative comparison, we evaluated the Kuramoto order parameter *R* as a measure of network synchrony (Fig. 5a). As shown previously, both the population bursting and sustained oscillation regimes are characterized by high synchrony in the network. We therefore superimposed the heatmap of *R* for the spiking network on the bifurcation diagram of the mean-field reduction over the same parameter range.

Within the parameter range examined here, the subcritical Andronov-Hopf and the inferred FLC curves lie close to one another and delimit a narrow bistable region, at the border between population bursting and asynchronous dynamics (both for the high and low activity side). For this reason, we highlight only the Andronov-Hopf bifurcation curve for clarity.

This analysis shows that the mean-field reduction effectively reproduces the qualitative dynamical behavior of the spiking network across a wide parameter range. Consistent with multiple previous studies [7, 34, 9], discrepancies between the two representations are concentrated at the boundaries between distinct dynamical regimes, where the mean-field reduction is less reliable.

In fact, as observed in the single-parameter analysis above, there is a mismatched between the two representations over the whole parameter range explored at the transition between sustained oscillation and the asynchronous regular regime. We notice that the mismatch is also consistent at the boundary between population bursting and low-activity asynchronous regime (rightmost Andronov-Hopf curve). Moreover, in this case, the increasing relevance of finite-size effects, highlighted in the previous section, manifests as a spatially fragmented and less sharply defined boundary within the network.

The boundary between population bursting and high-activity asynchronous regular regime is instead well captured by the mean field. This better agreement arises because the jump in mean firing rate (and, similarly, in mean membrane potential) is considerably sharper at this transition, consistently across the different values of *c*_*dopa*_ considered (Fig. 5b). Hence, the transition in the network is well captured by the model (dashed lines represent the *u*_*jump*_ values of the subcritical Andronov-Hopf bifurcation for the different values of*c*_*dopa*_).

The close correspondence at this transition may reflect the fact that the mean-field description is obtained using a first-order moment closure assumption and retains only a limited set of low-dimensional population observables, particularly the population firing rate and mean membrane potential. Higher-order fluctuations and the full microscopic distribution of neuronal states are not explicitly represented. Consequently, transitions dominated by finite-size fluctuations or changes in higher-order distributional structure are more difficult to reproduce precisely.

We further highlight that the mean field correctly reproduces the modulatory effect of dopamine, via the dopaminergic projection *c*_*dopa*_, on the activity of the neural population. Increasing *c*_*dopa*_ shifts the population from a low-activity asynchronous regime toward a regime of collective oscillatory or bursting activity and vice-versa. This modulatory effect is reproduced qualitatively by both the mean-field model and the spiking network.

To complement the qualitative comparison, we computed two additional metrics: the normalized Wasserstein distance and the Hellinger distance. The Wasserstein distance quantifies differences between the empirical distributions of population rate values obtained by flattening the time series over time. It therefore measures the similarity of the long-term marginal activity statistics, including quantities such as the mean activity and the range of the oscillations amplitude. The Hellinger distance, instead, quantifies differences between the normalized power spectra and therefore measures discrepancies in the distribution of spectral power across frequencies (Fig. 5c–d).

In the asynchronous regimes, the mean-field solution approaches a constant fixed point and therefore exhibits no appreciable power at nonzero frequencies. In the finite spiking network, by contrast, fluctuations persist around this value as a consequence of discrete spike emission. This is consistent with the deterministic mean-field description capturing the average population activity while neglecting fluctuations arising from the finite number of neurons. Consequently, the asynchronous regimes are characterized by low Wasserstein distance but high Hellinger distance.

Similarly, in the sustained oscillation regime, the low Wasserstein distance indicates that the two representations generate similar distributions of population rate values, whereas the larger Hellinger distance indicates a discrepancy in their spectral content. In this regime, the mean-field and network time series exhibit comparable oscillation amplitudes, while their dominant spectral peaks occur at different frequencies. Such a frequency mismatch can therefore result in a relatively large Hellinger distance even when the marginal distributions of population rate values remain similar.

The population bursting regime shows the strongest overall agreement according to the two metrics. Nevertheless, a mismatch becomes more pronounced in the vicinity of the transition associated with the rightmost Andronov-Hopf curve. In this region, the emergence of sparse population bursts leads to a disruption of the agreement in spectral content, particularly at low frequencies, between the mean-field and the neural population. This discrepancy is less apparent in the Wasserstein distance, since flattening the time series over time removes the temporal ordering of activity values and therefore discards information about the timing and inter-burst intervals. In contrast, the Hellinger distance retains sensitivity to these temporal differences through their effect on the power spectrum. The complementary nature of the two measures therefore provides a more complete characterization of the mismatch between the mean-field and spiking network representations.

Overall, although the two representations may differ in phase and do not overlap point by point in time, they can exhibit similar long-term marginal activity statistics within the same dynamical regime, as reflected by the Wasserstein distance. Their spectral agreement is more variable. In particular, the dominant bursting frequency is reproduced qualitatively, but even a moderate frequency shift can lead to a substantial increase in Hellinger distance.

### 2.4 Comparison of phase-amplitude coupling

In the previous section we showed that the best agreement between the mean-field and spiking network representations occurs within the region characterized by bursting dynamics. Because the bursting regime contains two distinct frequency components, we quantify phase-amplitude coupling (PAC) between the slow inter-burst rhythm and the fast intraburst oscillation. Specifically, we examine whether the phase of the slow burst repetition rhythm modulates the amplitude of the high-frequency intra-burst activity. The PAC is evaluated both in the mean-field description and in the spiking network by using the Tort Modulation Index (MI) [36, 37].

The results of this computation are shown in Fig. 6a, where the heatmap represents the computed MI for the mean-field and spiking network representations. We observe that both representations show similar overall values and a similar pattern within the parameter region of interest. MI is lower for small values of *u*_*jump*_ and gradually increases with it. Moreover, examining columns of the heatmap, MI tends to decrease as dopaminergic input increases, although this trend is less pronounced than the one observed for the adaptation parameter.

**Figure 6:**
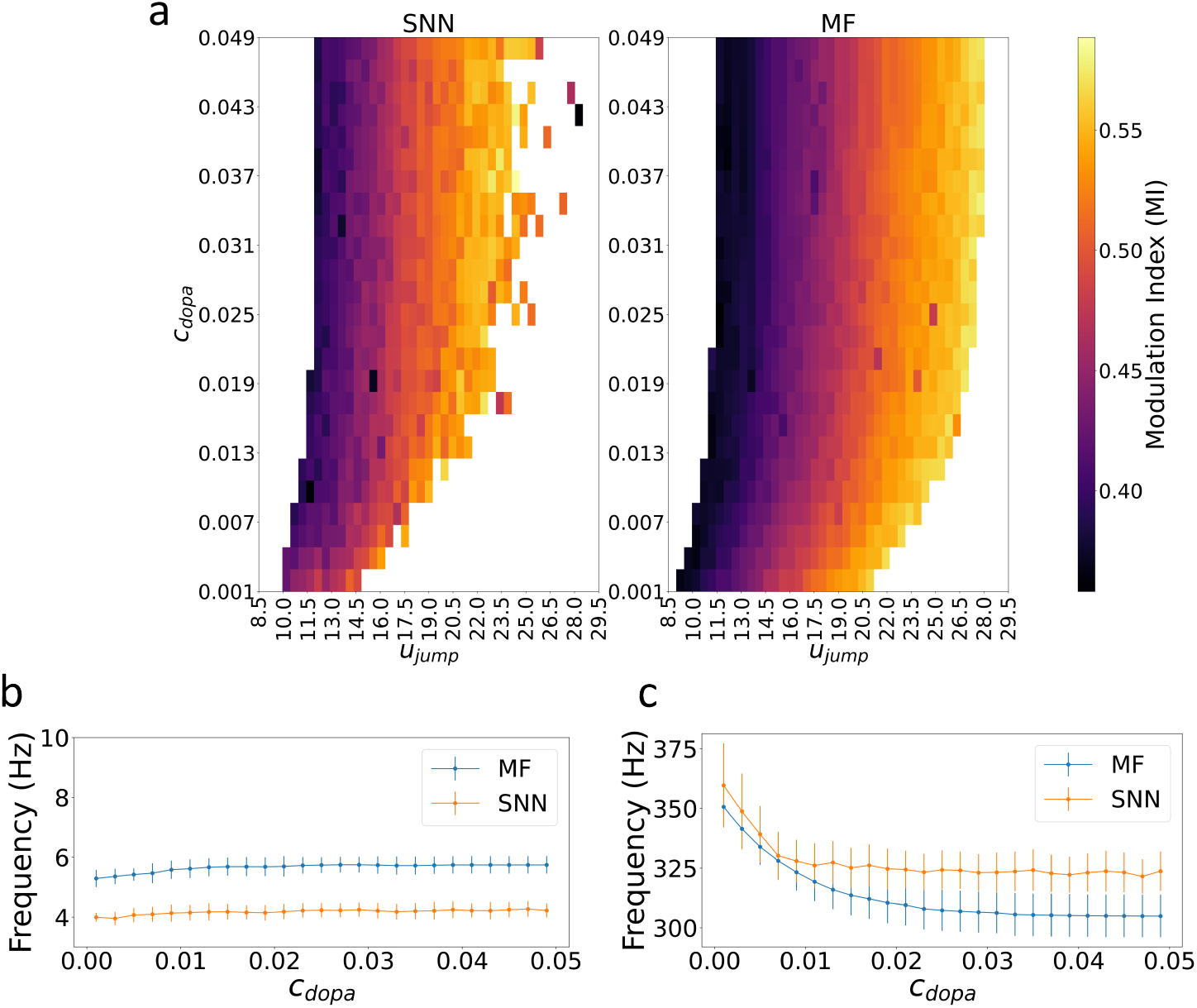
Phase-amplitude coupling between the slow and fast oscillatory components of bursting activity, in the spiking network (SNN) and mean-field (MF) representations. (a) Heatmap of the Tort Modulation Index (MI) across the (*u*_*jump*_, *c*_*dopa*_) parameter space, for the SNN (left) and the MF (right). White regions correspond to parameter combinations outside the bursting regime, where PAC is not defined. In both representations, MI is lowest near the onset of bursting (low *u*_*jump*_) and increases progressively with *u*_*jump*_, while showing a comparatively weaker decreasing trend with *c*_*dopa*_. The overall pattern is similar between the two representations, although the SNN estimate is visibly noisier, particularly at high *u*_*jump*_, reflecting the emergence of sparse burst events which affect the frequency content, especially regarding the slow inter-burst events. (b) Slow inter-burst frequency and (c) intra-burst HFO frequency, both as a function of *c*_*dopa*_, averaged over *u*_*jump*_ values. Blue line represents the MF and orange one the SNN. Errors bars indicate the standard deviation across the *u*_*jump*_ values within the bursting regime for each *c*_*dopa*_. In (b), both frequencies remain approximately constant across the explored range of *c*_*dopa*_ and fall within the 4–6 Hz band, with the MF systematically predicting higher values (∼5.2–5.7 Hz) than the SNN (∼4.0–4.3 Hz). In (c), both frequencies decrease with *c*_*dopa*_, more steeply and consistently in the MF, which continues to decrease across the full range explored, than in the SNN, which plateaus around *c*_*dopa*_ ≈ 0.02. The resulting offset between the two representations remains limited to approximately 20–40 Hz.

Examining the slow and fast frequency components around which the MI was computed, as functions of *c*_*dopa*_ (Fig. 6b–c, respectively), we find that both frequencies follow qualitatively similar trends in the spiking network and the mean-field reduction, and remain within overlapping ranges across the explored parameter space. The slow-oscillation frequency stays within the 4–6 Hz range in both representations, although the mean-field tends to predict systematically higher values than the spiking network. Similarly, while the HFO frequency decreases with *c*_*dopa*_ in both representations, the mean-field estimate falls increasingly below that of the network as *c*_*dopa*_ increases.

These results provide preliminary evidence that the mean-field reduction reproduces the presence and approximate strength of phase-amplitude coupling observed in the spiking network. However, a more complete analysis should examine the dependence of the MI on burst duration and inter-burst variability and the offset observed between the slow and fast frequency components of the two representations (Supplementary Material–Section B).

These results are intended to illustrate the mean-field model ability to capture spiking network behavior even for more complex features such as PAC, and should be regarded as preliminary. Achieving closer quantitative agreement in terms of frequency content, and more generally obtaining results directly comparable with experimental data, would require a more focused study of the precise parametrization of the two formulations across specific parameter regimes. This lies beyond the scope of the present work, whose contribution is primarily to explore whether distinct dynamical features of the spiking network can be captured by the mean-field reduction. Such further work would also help establish whether the frequency offset identified above reflects a genuine limitation of the mean-field reduction or an artifact of the specific parameter range explored here.

## 3 Discussion

We focus on a previously proposed mean-field reduction for a network of spiking Izhike-vich neurons [14], incorporating the modulatory effects of dopamine (DA) via activation of its specific D1-type receptor subtypes. We perform a systematic comparison between the reduced formulation and the underlying spiking network to understand in detail which aspects of the network dynamics are best captured by the mean field, both in terms of dynamical repertoire and of extracted features. In particular, we extend the work of Depannemaecker et al. [14] by systematically comparing the reduced formulation with the underlying spiking network and by quantifying which macroscopic observables and dynamical features are preserved by the reduction.

Through systematic bifurcation analysis, we identify multiple parameter domains, corresponding to qualitatively distinct dynamical regimes, namely sustained oscillations, population bursting, and an asynchronous regime. Because our reduction is derived directly from a population of spiking Izhikevich neurons, it preserves an explicit link to the underlying network through the description of collective variables, allowing us to validate the mean-field approximation directly against spiking network simulations under different conditions. Together with this qualitative validation, we also performed a systematic analysis of the differences between the two representations. We found good qualitative agreement between the two representations over a broad parameter range, as already highlighted in previous literature [9]. According to the selected distributional and spectral metrics, the closest correspondence occurs in the population bursting, although systematic differences in inter-burst timing remain, and asynchronous regime.

Nonetheless, our formulation has limitations, related to the specific methodological derivation of the mean-field reduction, which have already been extensively discussed [34, 9]. Several assumptions of the derivation (e.g., Lorentzian parameter distributions, moment-closure approximations) could lead to quantitative mismatches between the two formulations, which may be particularly relevant near critical parameter values or in highly heterogeneous networks. Moreover, in order to obtain a low-dimensional representation, conditions which are not biophysically plausible (infinite threshold and reset voltage, globally coupled network) were considered. Relaxing these assumptions could improve the quantitative accordance between the mean-field reduction and the spiking neural network together with an improvement of the description of microscopic mechanisms, as already partially proposed in some recent works [9, 6].

Finite-size effects represent an additional source of mismatch, which becomes relevant when reduced network activity coincides with proximity to a transition between dynamical regimes. This is exemplified by the end of the population-bursting regime (Fig. 4), where the regular burst recurrence predicted by the mean-field model gives way, in the spiking network, to long and irregular silent periods driven by fluctuations of the finite population, an effect that shifts and weakens as the network size increases.

Beyond this general validation, the population bursting regime is also of particular biological interest, since it is characterized by the co-occurrence of oscillations in a slow frequency band and in the high-frequency oscillations (HFOs) range, as already noted in the previous study [14].

HFOs comprise a broad class of electrophysiological oscillations, distinguished by their duration, frequency, and amplitude, and typically spanning a wide frequency range of approximately 60–1000 Hz [38]. They have been observed in various cortical and subcortical areas, both during normal processing [39, 38] and in pathophysiological conditions such as Parkinson’s disease [40, 41].

Hippocampal ripples (≈ 80–250 Hz) are a well-studied example of this kind of oscillation, and have been proposed to play an important role in memory consolidation [38]. In the context of PD, HFOs in the range of approximately 200–500 Hz, modulated by dopaminergic medication, have been observed in multiple studies [40, 42, 41].

Beyond the mere presence of HFOs, their coupling with lower frequencies has also been reported in the healthy brain, for example between hippocampal ripples and cortical sleep spindles or slow (delta) oscillations during memory consolidation [39, 38], and has been implicated in disease conditions. In Parkinsonian patients, dopaminergic medication reduces the coupling between HFOs and beta-band oscillations that is present in the untreated state, with coupling strength OFF medication reported to be up to two orders of magnitude greater than ON medication in the original report of this phenomenon [40].

We show that the current mean-field model is able to reproduce the coupling between slow and HFOs, as evaluated through the Modulation Index, both in terms of overall magnitude and of pattern across the parameter space. Hence, it is suited to describe and model mechanisms of frequency coupling as the one described above.

It has to be noted that the current work demonstrates the overall ability of the mean-field formulation to capture the dynamics of the spiking network, without the precise, situation-dependent parametrization that would be needed to adapt the system to the study of specific cases. However, because our reduction retains an explicit link to biophysically interpretable quantities, this limitation also points to a natural path forward: such parameters can, in principle, be estimated from experimental measurements and future work could fit them directly to experimental recordings, improving agreement with physiological conditions and enabling region-specific implementations. For example, PET-derived receptor densities could be used to obtain prior knowledge on receptor-specific parameters [43].

Moreover, given the flexibility in reproducing distinct dynamical regimes, the present mean-field model can further serve as a modular building block for whole-brain virtual twin models [44, 45, 46]. In this framework, different brain regions can be described by one or more nodes, each evolving according to a mean-field model incorporating region-specific neuromodulator profiles. By fitting regional model parameters to individual tractography and fMRI data, the current mean-field derivation could provide a basis for personalized simulations. Such personalized whole-brain models have shown promise in guiding clinical interventions in epilepsy [26, 47] and, with the incorporation of neuromodulatory dynamics, could likewise inform treatment strategies for disorders rooted in neuromodulatory dysfunction, such as Parkinson’s disease and schizophrenia [27, 28].

In this whole-brain context, our current formulation’s assumption of unidirectional coupling from the neuromodulator to the neural system could be relaxed by embedding reciprocal projections between nodes, where cortical regions project to dopaminergic nuclei, as suggested by [27] for DA, or by [43] in the context of serotonin modulation.

More generally, the mean-field framework we propose is not limited to reproducing DA-induced modulation of population activity through its receptor kinetics: it can be extended to other neuromodulators, both in the case of isolated modulatory substances and of combined regimes, by substituting or combining receptor-specific coupling functions. Prior research has underscored the critical role of neuromodulators, such as dopamine, serotonin, acetylcholine, and oxytocin, in enabling the brain’s diverse functional repertoire, and has shown that disruptions in neuromodulatory systems are associated with various pathological states [48, 43].

Taken together, these results show that the proposed mean-field reduction is capable of reproduce distinct dynamics of the underlying spiking network over a large region of the parameter space. By embedding receptor-mediated neuromodulation into an exact mean-field reduction of spiking networks, we thus provide a tractable yet biophysically interpretable framework for investigating how neuromodulators shape brain dynamics across scales, from single-neuron mechanisms to large-scale emergent behavior, offering a new computational tool for both basic research and clinical applications.

## 4 Methods

### 4.1 Single neuron model and mean-field reduction

We model each neuron as an adaptive quadratic integrate-and-fire (aQIF) unit ([49]). This model is widely used in the literature, as it captures the dynamical patterns of various neuron types through parameter variation. In the present work we extended the aQIF model with a dopaminergic modulation of synaptic efficacy following a recently published work ([14]). The membrane potential *v*_*i*_ and adaptation variable *u*_*i*_ of neuron *i* = 1, …, *N* evolve as

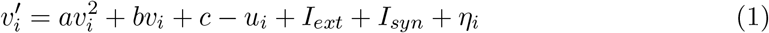

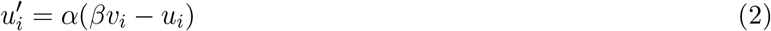

with the reset

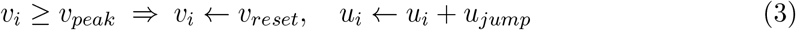

applied whenever the membrane potential reaches the threshold. The parameter *η*_*i*_ is a fixed random variable drawn from a distribution L(*η*) on (−∞, ∞), representing fixed variability in intrinsic excitability across the population. *I*_*ext*_ is a common external drive and *I*_*syn*_ the total synaptic input received from the network.

We consider an all-to-all connected population of excitatory neurons, coupled through AMPA-type synapses whose conductance is modulated by D1-type dopamine receptor activation, *M*_*D*1_:

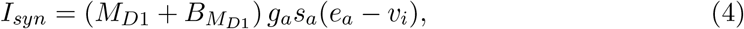

where *g*_*a*_ and *e*_*a*_ are the maximal conductance and reversal potential of the AMPA synapse, 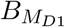 is the basal conductance offset, and *s*_*a*_ is the synaptic activation variable. Because coupling is all-to-all, *s*_*a*_ is homogeneous across the population and evolves according to

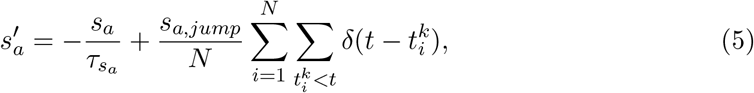

where 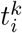 denotes the time of the *k*-th spike of neuron *i*.

The D1-receptor modulation *M*_*D*1_ relaxes exponentially toward a sigmoidal function of the extracellular dopamine concentration [*D*_*p*_] ([50]):

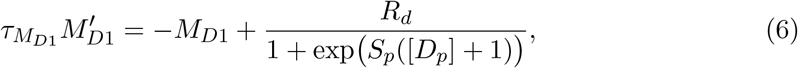

where *R*_*d*_ is the receptor density and *S*_*p*_ the receptor sensitivity to dopamine, with *S*_*p*_ *>* 0 (*S*_*p*_ *<* 0) corresponding to receptor activation (inhibition) by dopamine. Dopamine concentration itself follows Michaelis–Menten kinetics,

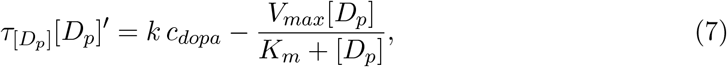

with *c*_*dopa*_ the input from dopaminergic afferents, scaled by a region-dependent factor *k*. We refer to ([14]) for the full motivation and characterization of this neuromodulation module. Since [*D*_*p*_] and *M*_*D*1_ evolve on time scales much slower than the firing-rate dynamics 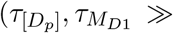 synaptic/firing time scales), we treat them as slow, quasi-static inputs that do not enter the mean-field reduction below.

We derive a low-dimensional mean-field approximation of the network dynamics in the limit *N* → ∞, following the approach of [34]. We summarize the main steps here. The complete derivation is given in the Supplementary Material–Sections C.

For large *N*, the network state is described by a population density *ρ*(*t, v, u, η*) over the single-neuron phase space (*v, u*) and heterogeneity *η*, obeying the continuity equation

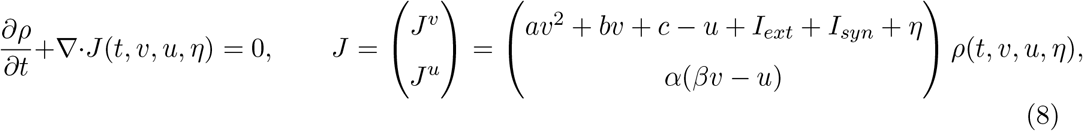

with boundary condition *J*^*v*^(*v*_*peak*_, *u*) = *J*^*v*^(*v*_*reset*_, *u*_*jump*_ + *u*) enforcing the reset (3). At threshold, the flux *J*^*v*^ equals the instantaneous population firing rate ([51]), which relates the synaptic and adaptation dynamics to *r*(*t*):

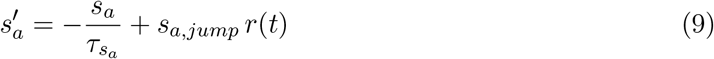

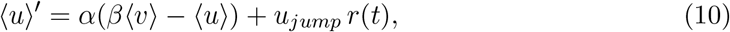

Under the assumption of weak spike-frequency adaptation (*u* ≫ *u*_*jump*_), so that fluctuations of individual *u*_*i*_ about ⟨*u*⟩ are negligible, we can derive a simplified equation for ⟨*u*(*t*)⟩ (a rigorous derivation from the continuity equation is given in Supplementary Material– Section C).

To close the system, we decompose *ρ*(*v, u, η*) = *ρ*^*u*^(*u* | *v, η*) *ρ*^*v*^(*v* | *η*) L(*η*) and apply the first-order moment closure ⟨*u* | *v, η*⟩ = ⟨*u* | *η*⟩ ([52, 53]), i.e. adaptation responds to the population-averaged membrane potential ⟨*v*(*t*)⟩ rather than to individual *v*_*i*_. Integrating the continuity equation over *u* then yields a reduced equation for *ρ*^*v*^(*t, v* | *η*) alone. Its stationary solution is Lorentzian in *v*, we assume that it follows a distribution of the same functional form in time ([7, 54]):

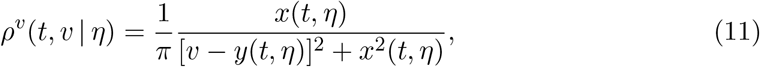

with *x* and *y* the half-width at half-maximum and center of the distribution. This yields

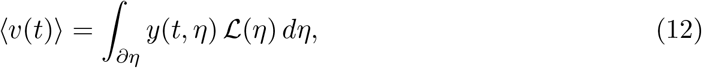

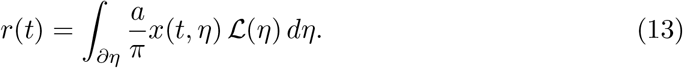

Although the infinite threshold and reset assumption is not plausible from a biological point of view, it is essential for the validity of the Lorentzian ansatz. In fact, it allows the passage from QIF model to theta-neuron model ([55]), via the change of variable *v*(*t*) = *tan*(*θ/*2). In the latter case, the ansatz we made corresponds to the OA ansatz ([54, 7]), used to investigate low-dimensional dynamics of a large population.

Substituting the ansatz (11) into the reduced continuity equation and requiring the resulting expression to vanish identically in *v* gives closed ODEs for *x*(*t, η*) and *y*(*t, η*).

Assuming, as is standard, that L(*η*) is itself Lorentzian,

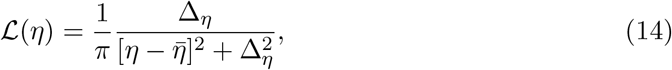

the integrals (12)–(13) can be evaluated in closed form via the residue theorem, yielding the mean-field system

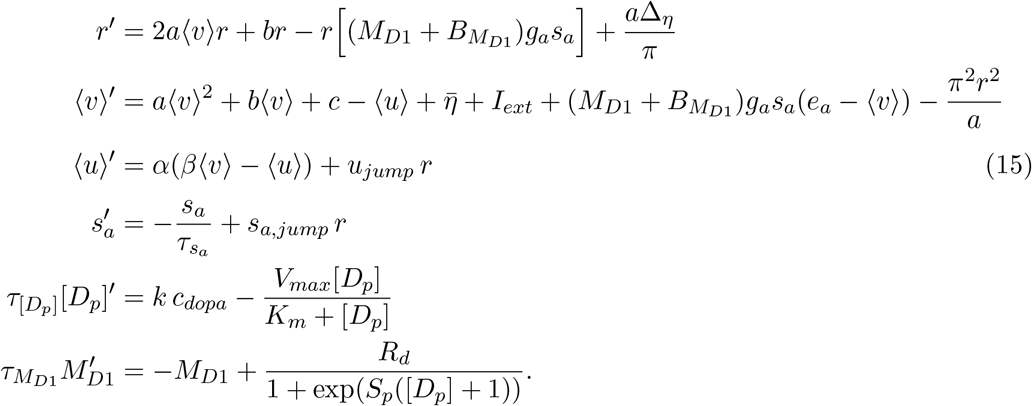

Numerical continuation and bifurcation analysis of the mean-field reduction (15) were performed in MATLAB using MatCont ([56, 57]).

### 4.2 Spiking network numerical simulations and analysis

For the spiking network, we simulated *N* = 5000 all-to-all coupled aQIF neurons as described in Section 4.1, using the Python package Brian2 ([58]). Simulation duration and time step depended on the feature being measured. In all cases we simulated at least a 5 s transient (increased close to transition or for specific analysis performed), which was discarded, and then recorded both the spiking activity of every neuron and the population firing rate. Heterogeneity was introduced by sampling the excitability parameter *η*_*i*_ deterministically from a Lorentzian distribution with mean 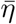 and half-width Δ_*η*_, using a quantile-based scheme to avoid the sampling artifacts of the unbounded Lorentzian tails at finite *N*. Parameter values for both the spiking network and the mean-field simulations are given in Table 1. Note that we used large, biologically unrealistic positive and negative values for *v*_*peak*_ and *v*_*reset*_ to approximate the theta-neuron limit underlying the Lorentzian ansatz (Section 4.1).

**Table 1:** Parameter values used in the spiking network and mean-field simulations. Unless otherwise stated, these values constitute the standard parameter set used throughout the study. The parameters *u*_*jump*_ and *c*_*dopa*_ are varied across simulations, with their specific values indicated in the corresponding figure captions.

|  |  |  |  |  |  |
| --- | --- | --- | --- | --- | --- |
| $a$ | 0.03 | $I_{ext}$ | 0 | $k$ | 1000 |
| $b$ | 5 | $g_a$ | 12 | $K_m$ | 150 |
| $c$ | 130 | $e_a$ | 0 | $V_{max}$ | 1300 |
| $\alpha$ | 0.013 | $s_{a,jump}$ | 0.8 | $R_d$ | 1 |
| $\beta$ | 0.4 | $\tau_{s_a}$ | 2.6 | $S_p$ | -1 |
| $u_{jump}$ | 12 | $\Delta_\eta$ | 5.0 | $c_{dopa}$ | 0.005 |
| $v_{peak}$ | 1000 | $\bar{\eta}$ | 10.0 | $\tau_{[D_p]}$ | 500 |
| $v_{reset}$ | -1000 | $B_{M_{D1}}$ | 0.2 | $\tau_{M_{D1}}$ | 500 |

We characterized the dynamical regime of the simulated network using different metrics. Global spike-timing variability was quantified with the coefficient of variation (CV) of the inter-spike intervals (ISIs). For neuron *i*,

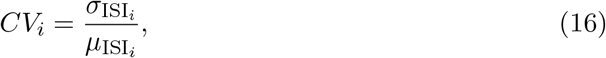

where 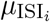 and 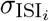 are the mean and standard deviation of neuron *i*’s ISIs. Values near zero indicate regular spiking, while values near or above unity indicate irregular, Poisson-like activity. However, the CV is sensitive to non-stationarities in the firing rate and to occasional extreme ISIs, which can dominate 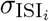 ([29, 30]).

For this reason we also used a complementary, local measure of variability, the CV2 ([31, 32]). For neuron *i* with *K*_*i*_ recorded ISIs, we first compute the local ratio for each pair of consecutive ISIs,

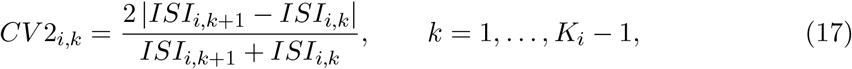

and then average over all pairs of neuron *i*’s ISIs to obtain a single value per neuron,

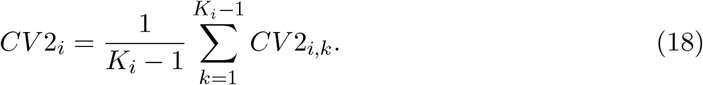

Because each term compares only two consecutive intervals rather than the full ISI distribution, the CV2 is more robust to rare extreme events (e.g., long inter-burst intervals in a population bursting regime) than the CV. As with the CV, values close to 0 indicate regular firing, values close to 1 correspond to irregular, Poisson-like activity, and values above unity indicate irregular, bursting-like local dynamics.

To evaluate network synchrony, we used the modulud of the Kuramoto order parameter ([33])

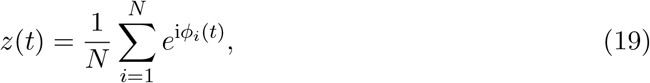

where *ϕ*_*i*_(*t*) is the instantaneous phase of neuron *i*, linearly interpolated between successive spikes:

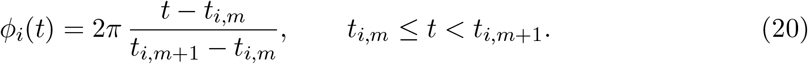

The modulus *R* = |*z*(*t*)| ranges from 0, indicating fully asynchronous firing, to 1, indicating that all neurons in the network fire in perfect synchrony.

To compare the long-term statistics of the mean-field and spiking network time series, we have flattened the values of each population-rate time series in time to obtain an empirical distribution. The resulting distributions are compared using the Wasserstein distance [59]. This measure quantifies the minimal cost of transforming one probability distribution into the other. For one-dimensional distributions it reduces to

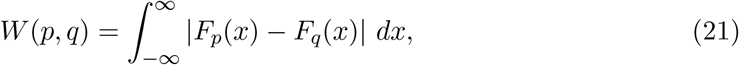

where *F*_*p*_(·) and *F*_*q*_(·) denote the cumulative distribution functions of *p* and *q*. The flattened distribution captures long-term statistical properties such as the mean activity level, variability, and the relative occupancy of low- and high-activity states, though it discards temporal ordering.

To obtain a more complete picture of the differences between the two systems, we additionally compared their spectral content using the Hellinger distance. This measure quantifies the overlap between corresponding bins of two distributions. It ranges from 0 (identical distributions) to 1 (disjoint, non-overlapping distributions). For discrete distributions it is defined as

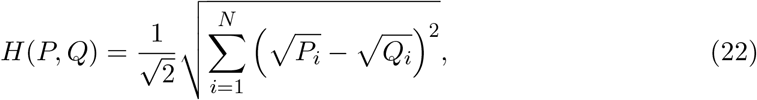

where *i* = 1, …, *N* indexes the frequency bins. The two spectra are computed on the same frequency grid, smoothed using the same procedure, and normalized before comparison, so that the Hellinger distance quantifies their overlap in spectral power and thereby captures differences in dominant frequencies and in the relative weight of slow and fast components. Wasserstein distance was computed using the specific function of the Python package SciPy, while Hellinger distance was manually implemented in Python.

### 4.3 Spectral features

To compute spectral features, we numerically simulated both the spiking network and the reduced mean-field system to obtain the population firing rate. Simulations were 30 s long with a time step of Δ*t* = 0.01 ms. From each simulation we discarded the first 10 s to avoid transient effects, as well as the final second, to avoid possible numerical errors at the simulation boundaries. Spectral analysis was performed with SciPy. The power spectral density was computed using Welch’s method with 2.7 s Hamming windows and 50% overlap.

To identify the frequency bands used to compute phase-amplitude coupling (PAC), we proceeded as follows. For the slow-oscillation band, the signal was first filtered in the range 0.5–50 Hz and smoothed with a Gaussian filter (*σ* = 2 Hz). This procedure was applied identically to the mean-field and spiking network signals. Peaks were then identified requiring a minimum separation of 3 Hz, and the highest peak was taken as the fundamental slow oscillation of the system. We defined the corresponding phase-frequency band as the 6 Hz-wide interval centered on this peak.

A similar procedure was used to identify the high-frequency oscillation (HFO) peak (100– 600 Hz), but with a larger Gaussian smoothing bandwidth (*σ* = 5 Hz), since the spectrum exhibited several closely spaced peaks in this range. We then defined the amplitude-frequency band of interest as the 100 Hz-wide interval centered on this peak.

Computation of PAC can be done by several means. Here we used the Modulation Index (MI) proposed by Tort et al. [36, 37], which quantifies coupling between a low-frequency phase component and a higher-frequency amplitude component. We chose this method because, when benchmarked against alternative approaches, it performed better in the presence of noise and was less sensitive to extreme amplitude values [60].

The signal was band-pass filtered separately within the phase-frequency band and the amplitude-frequency band defined above. The instantaneous phase and the amplitude envelope of the signal were obtained respectively from the angle and the magnitude of the analytic signal, computed via the Hilbert transform. The instantaneous phase values, ranging from −*π* to *π*, were divided into *N* = 20 equally spaced phase bins (18^°^ per bin). For each phase bin *i*, we computed the mean amplitude envelope across all samples whose instantaneous phase is within that bin, and normalized the resulting values to obtain a measure which behaves as a discrete probability distribution *P*:

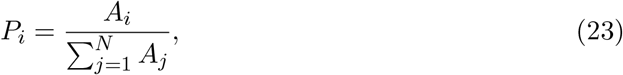

where *A*_*i*_ is the mean amplitude in phase bin *i*. The idea behind this measure is to quantify how much the distribution of these normalized amplitudes deviates from a uniform distribution *U*, which ideally corresponds to the absence of PAC.

Phase-amplitude coupling was quantified as the Kullback–Leibler divergence between the empirical and uniform distributions,

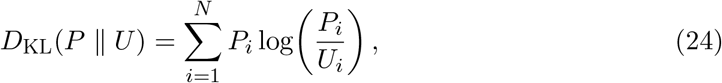

normalized by log(*N*) to obtain the Tort Modulation Index,

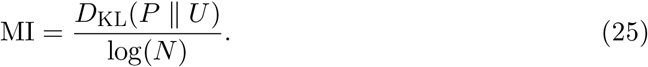

To avoid numerical issues from zero-valued probability bins, a small constant (*ϵ* = 10^−10^) was added to *P* before evaluating the logarithm. An MI of zero corresponds to a uniform amplitude distribution across phase bins, whereas higher MI values indicate an increasingly non-uniform, phase-dependent amplitude distribution, and therefore stronger phase-amplitude coupling.

## Author contribution

VJ, AS, ARM, SP, and PS were responsible for the conceptualization and for the funding acquisition. AF, DD and CD contributed to the conceptualization and research design, together with performing preliminary analysis. GC performed research and wrote the manuscript. DD supervised the work. All authors reviewed the manuscript. The authors have no conflict of interest to disclose.

## Acknowledgment

The project leading to this publication has received funding from the Excellence Initiative of Aix-Marseille Université - A*Midex, a French “Investissements d’Avenir programme” AMX-21-IET-017. This research has received funding from the European Union’s Horizon Europe Programme under the Specific Grant Agreement No. 101147319 (EBRAINS 2.0 Project). It has also received funding from the European Union’s Horizon Europe Programme under the Specific Grant Agreement No. 101137289 (Virtual Brain Twin Project), and No. 101057429 (project environMENTAL). This work has benefited from a government grant managed by the Agence Nationale de la Recherche (ANR) under the France 2030 program, reference ANR-22-PESN-0012. The authors acknowledge support from the grant “Personalised brain models of Parkinson’s disease patients”, “Modèle de cerveau personnalisé pour les patients atteints de la maladie de Parkinson”. Lithuanian-French programme Gilibert, project No. 2024-PRO-00148/ P-LZ-24-6, 2025-2026.

All authors declare no financial or non-financial competing interests.

## Code availability

The code associated to this article is available on GitHub at:

https://github.com/GabrieleCasagrande/Dopa_model

