## Supplementary Information for "Linking microscopic and mean-field descriptions of dopaminergic modulation in neural circuits"

### Supplementary material

#### Section A: Dynamical regimes of the spiking neural network

We report here the raster plots of the spiking neural network underlying the CV, CV2 and Kuramoto order parameter analyses presented in the main text (Figs. 2 and 3) for both transitions between dynamical regimes. Raster plots show $N = 1000$ neurons, randomly extracted from the $N = 5000$ neuron population used in the main text.

Figure 1 shows raster plots for the transition between sustained oscillations and the asynchronous regular state. At $u_{jump} = 6.0$ and $u_{jump} = 6.4$ (panels a, b), the network displays clear, regularly spaced vertical bands of synchronized activity, characteristic of the sustained-oscillation regime. As $u_{jump}$ increases toward 6.8 (panel c), these bands progressively dissolve into a spatially and temporally homogeneous pattern of activity, reflecting the loss of population synchrony and the emergence of the asynchronous regular state, consistent with the decrease in the Kuramoto order parameter $R$ reported in Fig. 2b of the main text.

Figure 2 shows raster plots for the transition between the asynchronous regular state and population bursting. Note the longer observation window (500 ms, compared to 100 ms in Fig. 1), chosen to accommodate the slower timescale of the bursting dynamics. At $u_{jump} = 9.513$ and $u_{jump} = 9.672$ (panels a, b), network activity remains spatially and temporally homogeneous, consistent with the asynchronous regular regime. At $u_{jump} = 9.882$ (panel c), the raster reveals the emergence of population-bursting events, visible as dense, synchronized vertical bands, separated by longer intervals of asynchronous activity, consistent with the sharp increase in the Kuramoto order parameter and the broadening of the CV distribution reported in Fig. 3a-b of the main text.

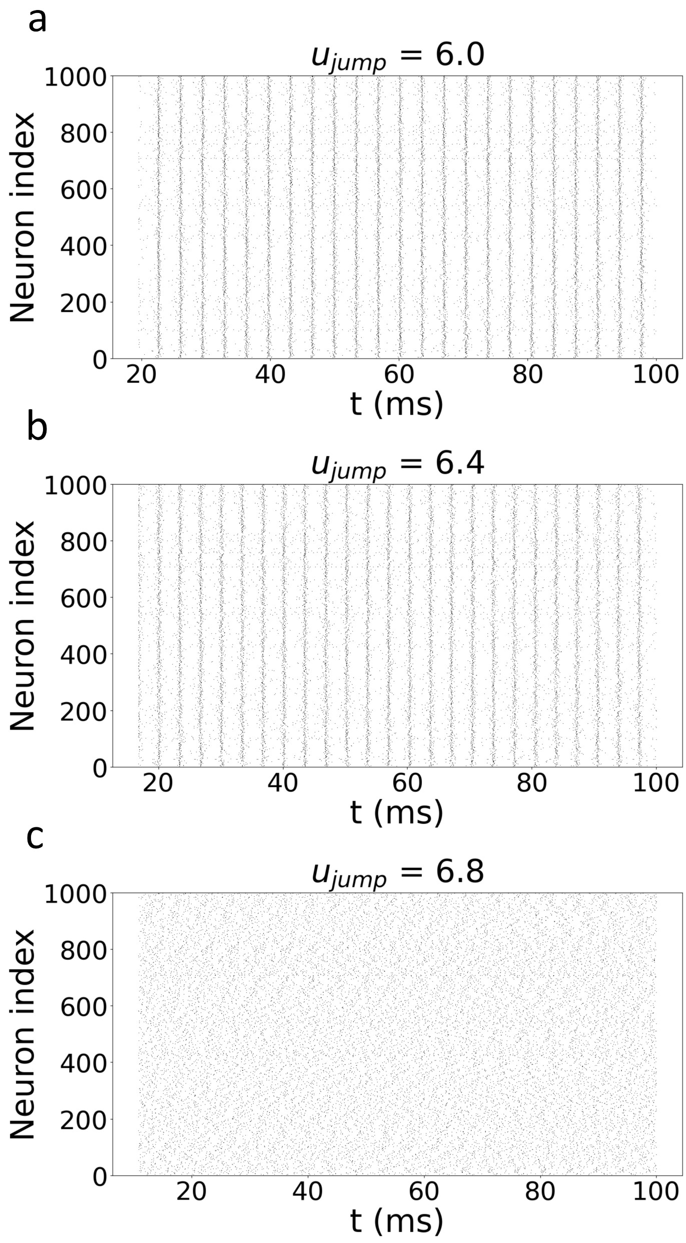


Figure 1: **Raster plots of the network activity across the transition between sustained oscillations and the asynchronous regular state.** Spiking activity of the network, shown for $u_{jump} = 6.0$ (a), $u_{jump} = 6.4$ (b), and $u_{jump} = 6.8$ (c), corresponding to the three values used for the CV and CV2 distributions in Fig. 2a of the main text. At $u_{jump} = 6.0$ and 6.4, regularly spaced vertical bands reflect synchronized, oscillatory population activity. At $u_{jump} = 6.8$, these bands are no longer visible, and neurons fire asynchronously while individually maintaining regular spiking, consistent with the asynchronous regular state described in the main text.

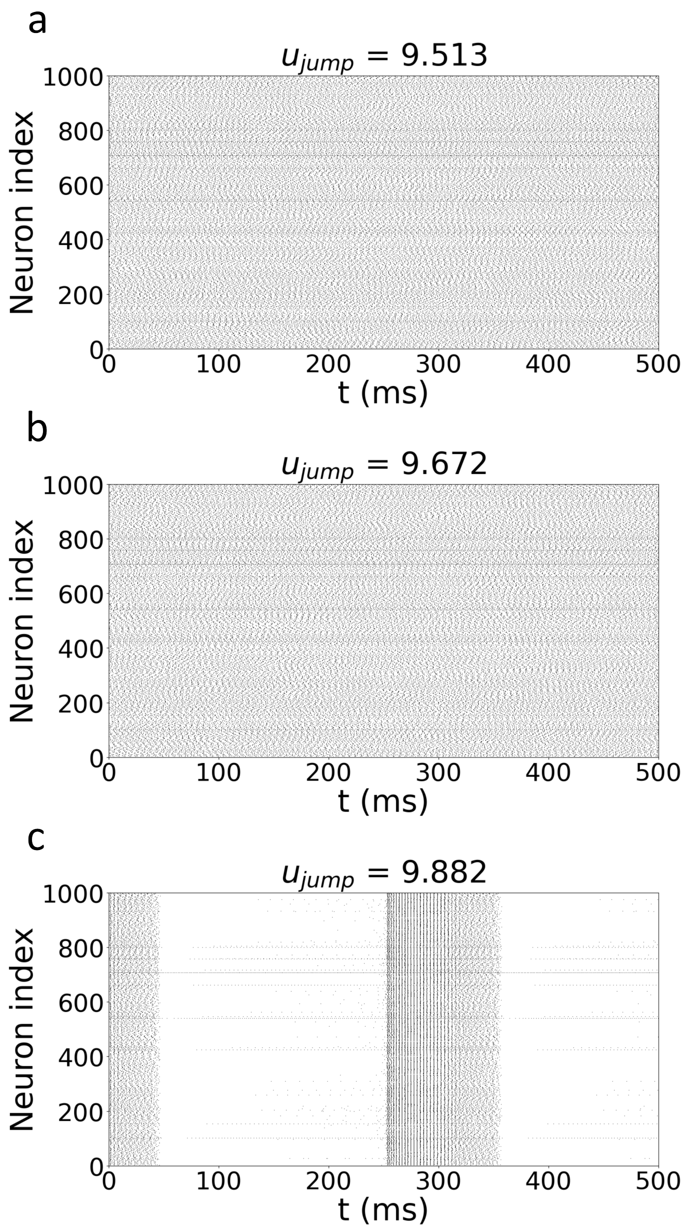


Figure 2: **Raster plots of the network activity across the transition between the asynchronous regular state and population bursting.** Spiking activity of the network, shown for $u_{jump} = 9.513$ (a), $u_{jump} = 9.672$ (b), and $u_{jump} = 9.882$ (c). At $u_{jump} = 9.513$ and 9.672, network activity remains homogeneous across neurons and time, characteristic of the asynchronous regular regime. At $u_{jump} = 9.882$, synchronized population-bursting events emerge (e.g., around $t \approx$ 0–50 ms and $t \approx$ 250–350 ms), separated by longer periods of quiescent activity, characterizing the microscopic dynamics of the bursting regime.

#### Section B: Analysis of the frequency content

As discussed in the main text, phase-amplitude coupling within the bursting regime involves two distinct frequency components: a slow rhythm associated with burst repetition, and a fast, high-frequency oscillation (HFO) associated with intra-burst spiking. While Fig. 6b–c of the main text report these two frequencies as a function of $c_{dopa}$ alone, Fig. 3 below provides their full two-parameter dependence, over the same $(u_{jump}, c_{dopa})$ range used in Fig. 6a, for both the spiking network (SNN) and the mean field (MF).

#### Section C: Details of mean-field derivation

In general, mean-field formalism captures the collective behavior of an ensemble of single units by describing the behavior of macroscopic observables. In the following, we consider specific averaged variables, which are suited to describe a population of spiking neurons:

- Mean membrane potential, defined as the average over the entire population of neurons:

$$\langle v(t)\rangle = \int_{\partial\eta}\int_{\partial u}\int_{\partial v} v\rho(t,v,u,\eta)d\eta du dv \quad (1)$$

- Mean adaptation current over the full network:

$$\langle u(t)\rangle = \int_{\partial\eta}\int_{\partial u}\int_{\partial v} u\rho(t,v,u,\eta)d\eta du dv \quad (2)$$

- Population firing rate, namely the flux of neurons through $v_{peak}$ over the whole range of $u$ and $\eta$:

$$r(t) = \int_{\partial\eta}\int_{\partial u} J^v(t,v_{peak},u,\eta,M_{D1},s)d\eta du \quad (3)$$

##### Spiking network model

The neuronal network from which we derive the mean-field reduction consists of microscopic units described by a specific parametrization of the adaptive quadratic integrate-and-fire (aQIF) model proposed by Izhikevich ([1]). This model is widely used in the

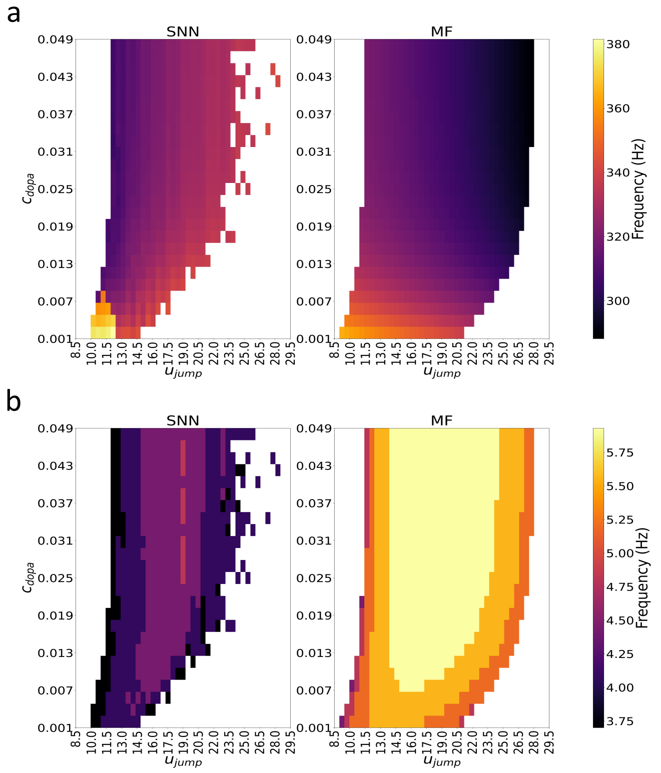


Figure 3: **Frequency content of the bursting regime across the** $(u_{jump}, c_{dopa})$ **parameter space, for the spiking network (SNN, left) and the mean field (MF, right).** (a) High-frequency oscillation (HFO) associated with intra-burst spiking activity, corresponding to Fig. 6c of the main text. Both representations show HFO frequencies in the $\sim$280–380 Hz range, with a similar overall trend along the $c_{dopa}$ axis: the frequency decreases as $c_{dopa}$ increases, although this decrease is sharper in the SNN than in the MF. Along the $u_{jump}$ axis, the two representations instead show an inverted gradient, with the MF frequency decreasing as $u_{jump}$ increases while the SNN frequency shows a slight increase. Despite this difference in gradient direction, the absolute discrepancy between the two representations remains limited, on the order of 20–40 Hz, a comparatively small offset given the overall HFO frequency range considered. As discussed in the main text, this analysis should be regarded as a proof of concept. A more precise parametrization would likely be needed to achieve closer quantitative agreement, although the overall trend in phase-amplitude coupling remains consistent between the two formulations. (b) Slow burst-repetition frequency, corresponding to Fig. 6b of the main text. In the SNN, this frequency remains largely confined to the lower half of the 4–6 Hz range explored ($\sim$4.0–4.5 Hz) across most of the parameter space. In the MF, the same frequency instead occupies mostly the upper half of this range ($\sim$5.25–5.75 Hz), with narrower bands at lower values near the edges of the bursting region. Both representations therefore remain within a broadly similar frequency band, though the MF appears to systematically predict a higher slow-oscillation frequency than the SNN. In both panels, white regions correspond to parameter combinations outside the bursting regime, where the corresponding frequency component is not defined.

literature, as it captures the dynamical patterns of various neuron types through parameter variation.

Each neuron is described by a two-dimensional system of ordinary differential equations governing the membrane potential v(t) and the adaptation current u(t):

$$v_i' = av_i^2 + bv_i + c - u_i + I_{ext} + I_{syn} + \eta_i \tag{4}$$

$$u_i' = \alpha(\beta v_i - u_i) \tag{5}$$

with $i = 1, 2, \ldots, N$.

When the membrane potential crosses the threshold, an action potential is generated and the following reset conditions apply:

$$\begin{aligned} v_i \geq v_{peak}, \text{ then } v_i \leftarrow v_{reset} \\ u_i \leftarrow u_i + u_{jump}. \end{aligned} \tag{6}$$

The parameter $\eta_i$ is a fixed random variable drawn from a distribution $\mathcal{L}(\eta)$ on $(-\infty, \infty)$. It represents a heterogeneous intrinsic current, different for all the neurons within the population. The term $I_{ext}$ denotes the common external input, while $I_{syn}$ represents the total synaptic input from other neurons in the network.

Neurons are connected through the following model for synaptic interactions, which takes into account modulation effect due to the activation of D1-type dopamine receptors, encoded in $M_{D1}$:

$$I_{syn} = (M_{D1} + B_{M_{D1}})g_a s_{a,i}(e_a - v_i) + g_g s_{g,i}(e_g - v_i), \tag{7}$$

where $g_{a,g}$ and $e_{a,g}$ are the maximum synaptic conductance and the reversal potential, respectively for the excitatory (AMPA) and inhibitory (GABA) synapses.

The variables $s_{a,i}$ and $s_{g,i}$ represent the synaptic activation, i.e. the portion of open ion channels in post-synaptic neurons due to the action of pre-synaptic ones. We assume the network to be all-to-all coupled, so that each neuron receives the summed output of all the pre-synaptic ones. As a consequence, both $s_{a,i}$ and $s_{g,i}$ are homogeneous across all the neurons. We refer at them as $s_a$ and $s_g, \forall i \in (1, N)$. In this work, we specifically focus on a network of excitatory all-to-all coupled neurons. Additional inputs, both excitatory and inhibitory, will be included when considering multiple interacting populations of neurons. The evolution equations for the synaptic activation reads:

$$\begin{cases} s'_a = -s_a/\tau_{s_a} + \frac{s_{j,a}}{N} \sum_{i=1}^{N} \sum_{t_i^k<t} \delta(t - t_i^k) \\ s'_g = -s_g/\tau_{s_g} \end{cases} \tag{8}$$

Here, $t_i^k$ is the time of the $k-th$ spike of neuron $i$ and $\delta(t)$ is the Dirac delta function. The modulation effect of dopamine receptors $M_{D1}$ presented in 7, is described by an exponential decrease and a sigmoid-shaped function [2], which represent the proportion of receptors activated as a function of the external dopamine concentration $[D_p]$:

$$\tau_{M_{D1}} M'_{D1} = -M_{D1} + \frac{R_d}{1 + exp(S_p([D_p] + 1))} \tag{9}$$

Here, $R_p$ represents the receptor density within a brain region, while $S_p$ corresponds to receptors sensitivity to variation in extracellular dopamine concentration. The sign of $S_p$ indicates whether the receptors will be activated ($S_p > 0$) or inhibited ($S_p < 0$) by the presence of dopamine.

As mentioned before, the term $[D_p]$ represents the concentration of dopamine molecules in the synaptic cleft, and its variation in time is described by:

$$\tau_{[D_p]}[D_p]' = kc_{dopa} - \frac{V_{max}[D_p]}{K_m + [D_p]} \tag{10}$$

The first term accounts for the increase due to the presence of projection from dopaminergic neurons, through $c_{dopa}$, scaled by a factor $k$ different for each region. The second term is the general form of the Michaelis-Menten equation [2]. For a detailed description of the terms accounting for dopamine modulation we refer to [3].

We point out that variation in dopamine concentration and its receptor activation is assumed to be much slower than the firing rate dynamics and therefore they do not affect the derivation of the mean-field system.

#### Mean-field reduction

In this section we summarize the mathematical steps to derive a low-dimensional mean-field model to approximate the behavior of a population of previously described neurons in the limit $N \to \infty$. The derivation is inspired by that presented in [4], so we briefly present the main ideas to retrieve the reduced set of equations. Complete derivation, comprising of all the extended passages, can be found in Supplementary Material.

Because of N is large, one can image that the evolution of the system variables describes a continuous flow in the space of variables. To describe the network, we consider a population density approach [5] by defining $\rho(t, v, u, s, \eta)$, which represents the density of neurons being, at time t, in $(v, u)$ position in the space of coordinates, with a certain value $\eta$ of the intrinsic current.

Assuming the total number of neurons to remain constant through the dynamics, the following continuity equation holds:

$$\frac{\partial \rho}{\partial t} + \nabla \cdot J(t, v, u, s, \eta) = 0 \tag{11}$$

Where $J(t, v, u, \eta)$ is the probability flux, defined as:

$$J = \begin{pmatrix} J^v \\ J^u \end{pmatrix} = \begin{pmatrix} G^v(v, u, \eta) \\ G^u(v, u) \end{pmatrix} \rho(t, v, u, \eta) = \begin{pmatrix} av^2 + bv + c - u + I_{ext} + I_{syn} + \eta \\ \alpha(\beta v - u) \end{pmatrix} \rho(t, v, u, \eta) \tag{12}$$

Taking into account the reset conditions (6), the following boundaries condition are defined:

$$J^v(v_{peak}, u) = J^v(v_{reset}, u_{jump} + u) \tag{13}$$

At the membrane potential threshold value, the flux can be seen as the instantaneous number of spikes fired by the neurons [6]. In this sense, the synaptic dynamics can be related to the population firing rate r(t), in the limit $N \to \infty$:

$$s_a' = -\frac{s_a}{\tau_{s_a}} + s_{a,jump} r(t), \tag{14}$$

with $s_{a,jump}$ and $\tau_{s_a}$ equal for each excitatory synapse (Supplementary Material - Section B).

Following the same reasoning, we can derive a simplified equation for $\langle u(t) \rangle$, if we consider small spike-frequency adaptation for all neuron, meaning $u >> u_{jump}$. Under this assumption, the dynamics of $\langle u(t) \rangle$ can be related to the r(t), namely:

$$\langle u(t) \rangle' = \alpha(\beta \langle v \rangle - \langle u \rangle) + u_{jump} \sum_{t^k < t} \delta(t - t^k) = \alpha(\beta \langle v \rangle - \langle u \rangle) + u_{jump} r(t) \tag{15}$$

This expression reflects the main idea of mean-field reduction itself, meaning that we individual fluctuation over single $u_i$'s are negligible.

Moving forward in the derivation, we decompose the probability density function in the following conditional form:

$$\rho(v, u, \eta) = \rho^u(u|v, \eta) \rho^v(v|\eta) \mathcal{L}(\eta) \tag{16}$$

The term $\mathcal{L}(\eta)$ is not time dependent but rather represent a "fixed" heterogeneity for the intrinsic current. As a consequence of the above decomposition, we can redefine $\langle v(t) \rangle$ and r(t). By integrating the continuity equation over the adaptation current $u$, we end up with a "reduced" continuity equation:

$$\frac{\partial}{\partial t} \rho^v(t, v|\eta) = -\frac{\partial}{\partial v} \left[ G^v(v, \langle u|\eta \rangle, \eta) \rho^v(t, v|\eta) \right] \tag{17}$$

In the last step, we exploit the first-order moment closure assumption $\langle u|v, \eta \rangle = \langle u|\eta \rangle$ [5, 7], which is fundamental to retrieve a closed set of equations. This means that $u_i$ is sensitive to $\langle v(t) \rangle$ rather than single values of $v_i$ [6, 8].

To further simplify mean-field equations, we proceed by making an ansatz for the functional form of $\rho^v(t, v|\eta)$. In particular, we notice that (17) has a Lorentzian shaped stationary solution. Thereby, we assume that it follows a distribution of the same functional form in time [9, 10], leading to:

$$\rho^v(t, v|\eta) = \frac{1}{\pi}\frac{x(t,\eta)}{[v - y(t,\eta)]^2 + x^2(t,\eta)}, \tag{18}$$

with $x(t, \eta)$ and $y(t, \eta)$ respectively the half-width at half-maximum and the location of the center of the distribution.

Taking into account this expression for the conditional density function and assuming $v_{peak} = -v_{reset} \to \infty$, one can express $\langle v(t)\rangle$ and r(t) in term of the Lorentzian distribution parameters:

$$\langle v(t)\rangle = \int_{\partial\eta} y(t,\eta)\mathcal{L}(\eta)d\eta \tag{19}$$

$$r(t) = \int_{\partial\eta} \frac{a}{\pi}x(t,\eta)\mathcal{L}(\eta)d\eta \tag{20}$$

Although the infinite threshold and reset assumption is not plausible from a biological point of view, it is essential for the validity of the Lorentzian ansatz. In fact, it allows the passage from QIF model to theta-neuron model [11], via the change of variable $v(t) = tan(\theta/2)$. In the latter case, the ansatz we made correspond to the OA ansatz [10, 9], used to investigate low-dimensional dynamics of large population.

Next, we proceed to plug (18) into the reduced continuity equation (17) to obtain the time evolution for $x(t, \eta)$ and $y(t, \eta)$. The result is a second order equation in $v$. To eliminate v-dependence (i.e. equation valid for each value of v), we consider the coefficients of $v^2$ and $v$ to be equal to 0. We find functional form for $x'(t, \eta)$ and $y'(t, \eta)$ and, once plugged into the function for the coefficient related to $v^0$ it disappears (Supplementary Material - Section D).

Up to this point, we have obtained the mean-field equations for $r(t, \eta)$ and $\langle v(t, \eta)\rangle$ as functions of the intrinsic current $\eta$. To retrieve the final set of mean-field equations, we need to choose a functional form for the heterogeneity distribution to compute the integrals (19) and (20). For simplicity, we assume that $\mathcal{L}(\eta)$ is also distributed according to a Lorentzian distribution [9, 4]:

$$\mathcal{L}(\eta) = \frac{1}{\pi} \frac{\Delta_\eta}{[\eta - \bar{\eta}]^2 + \Delta_\eta^2} \tag{21}$$

with $\Delta_\eta$ and $\bar{\eta}$ respectively the half-width at half-maximum and the centroid of the distribution.

We stress that the choice of $\mathcal{L}(\eta)$ is arbitrary. However, final results, obtained with different functional forms, are qualitatively similar to those obtained using the Lorentzian ansatz, as shown in [9].

Exploiting the residue theorem (Supplementary Material - Section E) we achieve the mean-field set of equations:

$$\begin{aligned}
r(t)' &= 2a\langle v\rangle r + br - r\left((M_{D1} + B_{M_{D1}})g_a s_a + g_g s_g\right) + \frac{a\Delta_\eta}{\pi} \\
\langle v\rangle' &= a\langle v\rangle^2 + b\langle v\rangle + c - \langle u\rangle + \bar{\eta} + I_{ext} + (M_{D1} + B_{M_{D1}})g_a s_a(e_a - \langle v\rangle) + g_g s_g(e_g - \langle v\rangle) - \frac{\pi^2 r^2}{a} \\
\langle u\rangle' &= \alpha(\beta\langle v\rangle - \langle u\rangle) + u_{jump} r(t) \\
s_a' &= -s_a/\tau_{s_a} + s_{a,jump} r(t) \\
s_g' &= -s_g/\tau_{s_g} \\
\tau_{[D_p]}[D_p]' &= kc_{dopa} - \frac{V_{max}[D_p]}{K_m + [D_p]} \\
\tau_{M_{D1}} M_{D1}' &= -M_{D1} + \frac{R_d}{1 + exp(S_p([D_p] + 1))}
\end{aligned} \tag{22}$$

#### Evolution of synaptic dynamics

Consider the single exponential synapse model we used for the excitatory synapse:

$$s_a' = -s_a/\tau_{s_a} + \frac{s_{j,a}}{N} \sum_{i=1}^{N} \sum_{t_i^k < t} \delta(t - t_i^k)$$

Here we demonstrate that, considering $N \to \infty$:

$$s'_a = -s_a/\tau_{s_a} + \frac{s_{j,a}}{N}\sum_{i=1}^{N}\sum_{t_i^k<t}\delta(t-t_i^k) = -s_a/\tau_{s_a} + s_{j,a}j(t) = -s_a/\tau_{s_a} + s_{j,a}r(t)$$

where we highlight the contribute:

$$j(t) = \frac{1}{N}\sum_{i=1}^{N}\sum_{t_i^k<t}\delta(t-t_i^k)$$

First, let us define $n_i(t)$ the number of spikes fired by the i-th neuron in $[0,t]$. Its average value across all the network will be:

$$\langle n_i(t)\rangle = \lim_{N\to\infty}\frac{1}{N}\sum_{i=0}^{N}\int_0^t\sum_{x_i^k<t}\delta(x-x_i^k)dx = \lim_{N\to\infty}\int_0^t j(x)dx$$

Next, we can define the population firing rate as:

$$r(t) = \lim_{\Delta t\to 0}\frac{1}{\Delta t}\lim_{N\to\infty}\frac{1}{N}\sum_{i=1}^{N}\frac{n_i(t+\Delta t)-n_i(t)}{N}$$

Finally, taking the limit $N \to \infty$ the previous equation can be rearranged as:

$$r(t) = \lim_{\Delta t\to 0}\frac{\langle n_i(t+\Delta t)\rangle - \langle n_i(t)\rangle}{\Delta t} = \frac{d}{dt}\langle n_i(t)\rangle$$

where the last term can be replaced by $j(t)$ in the thermodynamic limit, as shown before.

#### Derivation of $\langle u(t)\rangle$

In the following we will explain in detail the derivation of:

$$\langle u(t)\rangle = \int_{\partial\eta}\int_{\partial u}\int_{\partial v} u\rho(t,v,u,\eta)d\eta du dv$$

First of all, by exploiting the definition of the continuity equation we can separate the integral in 2 parts:

$$\langle u(t)\rangle' = \int_{\partial\eta}\int_{\partial u}\int_{\partial v} u\frac{\rho}{\partial t}d\eta dudv = -\int_{\partial\eta}\int_{\partial u}\int_{\partial v} u(\frac{\partial J^v}{\partial v} + \frac{\partial J^u}{\partial u})d\eta dudv$$

Consider the first one and proceed to integrate over $\partial v$:

$$\int u\left(\frac{\partial J^v}{\partial v}\right)dvdud\eta = \int uJ^v|_{\partial v}dud\eta \approx u_{jump}\int\left[(uJ^v)|_{\partial u} - \int J^v(v_{peak})dv\right]d\eta du =$$
$$= -u_{jump}\int J^v(v_{peak})dud\eta$$

In the second passage, we assume that $\langle u|\eta\rangle \gg u_{jump}$ and perform a Taylor expansion. Afterward, we use the fact that the flux disappears at the boundaries of $u$.
The second term instead reads:

$$\int u\left(\frac{\partial J^u}{\partial u}\right)dvdud\eta = \int\left[(uJ^v)|_{\partial u} - \int J^u du\right]dvd\eta = -\langle G^u\rangle = -\alpha\left(\beta\langle v\rangle - \langle u\rangle\right)$$

Where, in the second step, we consider once again the disappearance of the flux at the boundary of $u$, together with the definition of mean value, since $J^u = G^u\rho(v,u,\eta)$, and its linearity. Combining the two results the following expression for the evolution of the mean adaptation current is found:

$$\langle u(t)\rangle' = \alpha(\beta\langle v\rangle - \langle u\rangle) + u_{jump}r(t)$$

**Relation between mean-field variables and Lorentzian distribution parameters**

First, since Lorentzian distribution has mean only in principal value sense, $y(t)$ is defined in the following way:

$$y(t) = P.V.\int_{\partial v} v\rho^v(t,v|\eta)dv = \langle v(t,\eta)\rangle$$

The mean membrane potential can thus be considered as:

$$\langle v(t) \rangle = \int_{\partial\eta} y(t,\eta)\mathcal{L}(\eta)d\eta$$

Moreover, under the condition $v_{peak} = -v_{reset} \to \infty$, the population firing rate can be related to the Lorentzian distribution parameters:

$$r(t,\eta) = \lim_{v\to\infty} G^v(v, \langle u|\eta\rangle, \eta, M_{D1}, s)\rho^v(t, v|\eta) \propto a v_{peak}^2 \cdot \frac{x(t,\eta)}{\pi v_{peak}^2} = \frac{a}{\pi} x(t,\eta)$$

By integrating over the noise we get the population firing rate:

$$r(t) = \int_{\partial\eta} \frac{a}{\pi} x(t,\eta)\mathcal{L}(\eta)d\eta$$

Next we proceed to substitute the Lorentzian ansatz into the continuity equation for the probability density of mean membrane potential to obtain the time evolution for the parameters of the distribution. We obtain a second order equation in $v$ and equating the coefficients of $v^2$ and $v$ to 0, we find respectively:

$$x'(t,\eta) = 2axy + bx - x\left((M_{D1} + B_{M_{D1}})g_a s_a + g_g s_g\right)$$
$$y'(t,\eta) = ay^2 + by + c - ax^2 - \langle u|\eta\rangle + \eta + I_{ext} + (M_{D1} + B_{M_{D1}})g_a s_a(e_a - y) + g_g s_g(e_g - y)$$

Both of the results, lead to the disappearance of the coefficient related to $v^0$. By defining a new complex variable $z(t,\eta) = x(t,\eta) + iy(t,\eta) = \frac{\pi}{a} r(t,\eta) + i\langle v(t,\eta)\rangle$ we can write the reduced continuity equation in the form:

$$\frac{\partial}{\partial t} z = z\left(b - (M_{D1} + B_{M_{D1}})g_a s_a - g_g s_g\right) + i\left[-az^2 - \langle u|\eta\rangle + \eta + c + I_{ext} + (M_{D1} + B_{M_{D1}})g_a s_a e_a + g_g s_g e_g\right]$$

#### Computation of the final mean field equations using the residue theorem

In this section we further explain the computation of the integrals 19 and 20. Let us consider the integral:

$$\oint_{\Gamma_R} f(\eta)d\eta = \int_{-R}^{R} f(\eta)d\eta + \int_{\gamma_R} f(\eta)d\eta$$

where $\Gamma_R$ is an oriented rectifiable curve and $\gamma_R$ is the semicircle centered at the origin and connecting -R to R through the negative part of the imaginary plane.

Now assume:

$$\begin{aligned} f(\eta) &= \frac{a}{\pi}x(\eta,t)\cdot\mathcal{L}(\eta) = \frac{a}{\pi^2}x(\eta,t)\cdot\frac{\Delta_\eta}{[\eta-\bar{\eta}]^2+\Delta_\eta^2} = \\ &= \frac{a}{\pi^2}x(\eta,t)\cdot\frac{1}{2i}\left(\frac{A}{\eta-(\bar{\eta}-i\Delta_\eta)} - \frac{B}{\eta-(\bar{\eta}-i\Delta_\eta)}\right) = \frac{a}{\pi^2}x(\eta,t)\cdot\frac{1}{2i}\left(\frac{A}{\eta-\eta_1} - \frac{B}{\eta-\eta_2}\right) \end{aligned}$$

Using the residue theorem one would get:

$$\oint_{\Gamma_R} f(\eta)d\eta = -2\pi i\cdot Res_{\eta=\eta_2} f(\eta) = \frac{a}{\pi}x(\bar{\eta}-i\Delta_\eta,t)$$

To recover the result we are looking for we use the Jordan lemma to demonstrate that indeed $\int_{\gamma_R} f(\eta)d\eta \to 0$ as $R\to\infty$.

First, from the ML inequality we obtain:

$$\left|\int_{\gamma_R} f(\eta)d\eta\right| = \left|\int_{\gamma_R}\frac{a}{\pi^2}x(\eta,t)\frac{\Delta_\eta}{(\eta-\bar{\eta})^2+\Delta_\eta^2}d\eta\right| \le l_{\gamma_R}\cdot\sup_{\gamma_R}\left|\frac{a}{\pi^2}x(\eta,t)\frac{\Delta_\eta}{(\eta-\bar{\eta})^2+\Delta_\eta^2}\right|,$$

with $l_{\gamma_R} = \frac{1}{2}(2\pi R) = \pi R$.

Next, we exploit the triangular inequality to deal with the second part of the last equation:

$$(\eta-\bar{\eta})^2 = \left|(\eta-\bar{\eta})^2\right| = \left|(\eta-\bar{\eta})^2+\Delta_\eta^2-\Delta_\eta^2\right| \le \left[(\eta-\bar{\eta})^2-\Delta_\eta^2\right]+\Delta_\eta^2.$$

Using the above relation and letting $\eta \to \pm\infty$,i.e. when we extend the radius $R \to \infty$, we obtain:

$$\left|\frac{1}{(\eta-\bar{\eta})^2+\Delta_\eta^2}\right| \leq \frac{1}{|\eta-\bar{\eta}|-\Delta_\eta^2} = \frac{1}{R^2-\Delta_\eta^2}.$$

Moreover, we make the reasonable assumption that the half-width at half-maximum $x(\eta, t)$ is bounded by an upper value M. As a result:

$$\left|\frac{a}{\pi^2}x(\eta,t)\frac{\Delta_\eta}{(\eta-\bar{\eta})^2+\Delta_\eta^2}\right| \leq M\frac{a\Delta_\eta}{\pi^2}\frac{1}{R^2-\Delta_\eta^2}.$$

Finally, putting together all the previous results

$$\lim_{R\to+\infty} l_{\gamma_R}\cdot\sup_{\Gamma_R}\left|\frac{a}{\pi^2}x(\eta,t)\frac{\Delta_\eta}{(\eta-\bar{\eta})^2+\Delta_\eta^2}\right| = \lim_{R\to+\infty}\pi RM\frac{a\Delta_\eta}{\pi^2}\frac{1}{R^2-\Delta_\eta^2} = 0$$

This demonstrate that indeed that the integral computed over the semicircle $\gamma_R$ approaches 0 when $R \to \infty$, hence:

$$r(t) = \oint_{\Gamma_R} f(\eta)d\eta = \int_{-\infty}^{\infty} f(\eta)d\eta = \frac{a}{\pi}x(\bar{\eta}-i\Delta_\eta, t)$$

Exploiting a similar derivation we can also see that:

$$\langle v(t)\rangle = \int_{-\infty}^{\infty} y(\eta,t)\mathcal{L}(\eta)d\eta = y(\bar{\eta}-i\Delta_\eta, t)$$
